# Medial accumbens shell spiny projection neurons encode relative reward preference

**DOI:** 10.1101/2022.09.18.508426

**Authors:** Christian E. Pedersen, Raajaram Gowrishankar, Sean C. Piantadosi, Daniel C. Castro, Madelyn M. Gray, Zhe C. Zhou, Shane A. Kan, Patrick J. Murphy, Patrick R. O’Neill, Michael R. Bruchas

## Abstract

Medial nucleus accumbens shell (mNAcSh) is a critical brain region for driving motivated behaviors. Despite this well-established role, the underlying reward processing of individual neurons, circuits and cell-types within mNAcSh remains largely unknown. Here, we leverage deep brain 2-photon calcium imaging through endoscopic lenses to record mNAcSh spiny projection neuron (SPN) ensemble responses to rewards of different concentrations and to reward-predictive cues across cue-reward learning. Reward responses were found to be heterogeneous and particularly differentiated based on reward concentration and cell type. A large subpopulation of reward-excited enkephalinergic SPNs were found to be specifically recruited during consumption of high concentration, unpreferred reward. A major enkephalinergic efferent projection from mNAcSh to ventral pallidum (VP) was also found to be recruited to high concentration but unpreferred reward and to causally drive low positive reward preference. Enkephalin and dynorphinergic SPNs in mNAcSh distinctly represent rewards of different preference and propagate distinct signals through efferent projections to drive consummatory behavior.

---

The brain must compute and represent distinct identities of rewards to facilitate reward preference and to drive reward acquisition. The nucleus accumbens (NAc) is established as a major brain region for driving reward seeking and consumption ^1,2^. The NAc has many glutamatergic afferents such as basolateral amygdala, prefrontal cortex, ventral pallidum (VP) and ventral hippocampus and receives dopaminergic innervation from ventral tegmental area (VTA) ^3^. These NAc afferents have been shown to be highly reward-modulated, and in many cases, even reflect the quantity and preference of rewards ^4–9^.

The NAc is primarily composed of two dissociable types of spiny projection neurons (SPN) that are canonically defined by their expression of either dopamine 1 (D1R) or dopamine 2 (D2R) type receptors. D1R-SPNs also largely co-express the endogenous opioid peptide dynorphin, while D2R-SPNs primarily express enkephalin ^10,11^. Previous studies have shown NAc SPNs to be modulated by rewards and reward-predictive cues but have not resolved the molecular identity of the recorded neurons or tracked changes in activity of individual neurons across multiple days of cue-reward learning ^12,13^. Most reports have also primarily focused on NAc core and have not given specific attention to the reward-responsivity of the medial shell subregion (mNAcSh). mNAcSh has been shown to be very distinct from NAc core in its connectivity, function and dopaminergic innervation ^3,14–18^. Even within mNAcSh, functional distinctions have been isolated across anterior-posterior and dorsal-ventral axes ^2,11^.

Here, we sought to characterize the responses of individual mNAcSh SPNs to rewards of different identities and across cue-reward learning with particular focus on the molecular identity and relative anatomical position of the recorded cells. Recent experimental advances have allowed the simultaneous recording of dozens of genetically defined neurons in deep brain structures using 2-photon calcium imaging through endoscopic lenses ^19,20^. The low noise-floor of 2-photon microscopy also allows for the resolution of individual cell morphology and permits high fidelity tracking of individual neuron activity across multiple days during repeated training and behavioral sessions ^21^. In the current study, we leverage expertise in 2-photon imaging within deep brain to conduct longitudinal studies on how mNAcSh SPNs respond to multiple reward identities and change across cue-reward learning across time. This approach revealed a distinct subpopulation of enkephalinergic SPNs (Penk-SPN) that are selectively activated during the consumption of a less preferred reward. We then isolate a Penk-SPN circuit from mNAcSh to ventral pallidum (VP) as selectively activated during less preferred reward to causally drive a low, positive reward preference. These results provide foundational insight into how relative reward preference information is integrated and separated across mNAcSh neuronal ensembles and how these SPNs influence the drive for consummatory behavior.

## Results

### Differing concentrations of reward recruit distinct mNAcSh SPN subpopulations

Previous studies have consistently found that mice and rats prefer sucrose concentrations between 10-15% and that the distribution of sucrose preferences is shifted toward lower concentrations during water deprivation ^22–25^. Mice also tend to drink smaller volumes of higher concentration sucrose (>20%) due to satiety ^26^. In order to characterize differences in how rewards of varying preference are processed in mNAcSh, we presented mice with 4 distinct reward conditions that had differing concentration: 0% (water), 3%, 10% and 30% sucrose. To test the relative preference of the 4 sucrose reward conditions while minimizing effects from satiety, we tested mice in an adapted 2-bottle choice assay where the animals were only given brief access (20 s) to sucrose sippers with a variable intertrial interval between access periods. (**Fig. 1a; Extended Data Fig. 1a-b**). Every other day (odd days) both bottles contained 10% sucrose to control for side bias, with mice displaying no bias toward the left or right sipper on any day, suggesting that prior exposure to differing sucrose concentrations did not produce any bias (**Extended Data Fig. 1c-d**). Consistent with the aforementioned reports, we found that water-restricted C57 mice preferred 10% sucrose over all other conditions, and 30% sucrose less than all other conditions (**Fig. 1b**).

**Fig. 1.**
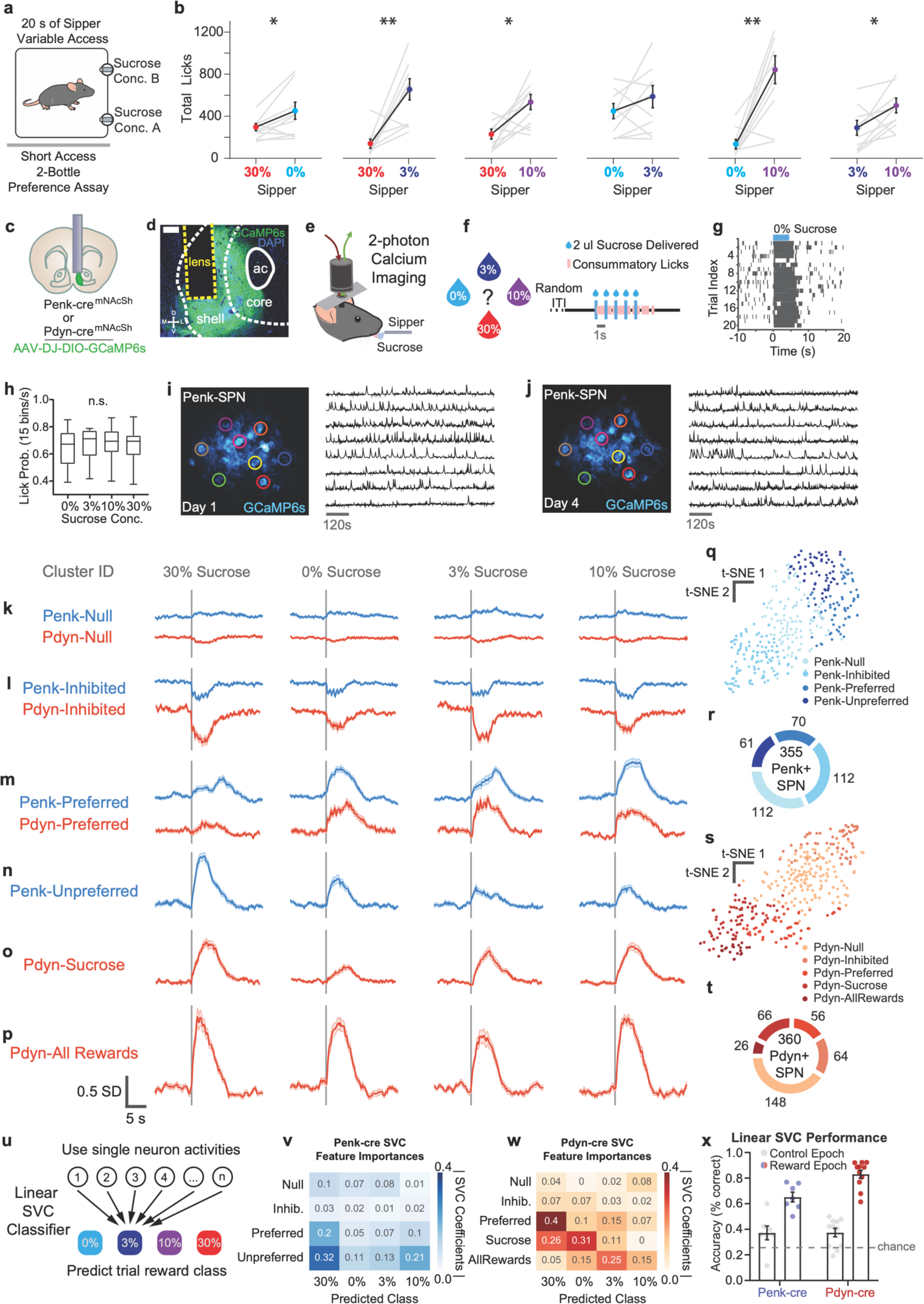
Distinct subpopulations of mNAcSh SPNs are recruited in response to rewards of different preference. (a) Schematic of brief access 2-bottle choice task: animal makes choice between two sippers containing different concentrations of sucrose reward. (b) Preference between each pairing of the 4 sucrose concentration conditions; graphs rearranged in order of preference (n = 10 animals, pairwise t-test: *p = 0.031, **p = 0.003, *p = 0.012, p = 0.227, **p = 0.001, p = *0.026). (c) Coronal section cartoon displaying target brain region, cell-types and lens implant location. (d) Schematic of 2-photon calcium imaging through endoscopic lens during consumption of variable sucrose reward. (e) Time-course of sucrose reward delivery for a given trial. Droplets were either 0%, 3%, 10% or 30% sucrose solution in water. (f) Representative lick raster depicting individual licks that were detected by contact licometer during behavior and imaging session. (g) Mean lick probability during reward epoch (0-8 seconds from reward onset) for each sucrose reward condition (n = 18 animals; one-way ANOVA: p = 0.9275). (h) t-sne plot of variance in activity of Penk-SPN clusters during all reward conditions. Each dot represents an individual tracked neuron that composes the cluster. (i) Fraction of all tracked Penk-SPN neurons that belong to each functional cluster. (j) t-sne plot of variance in activity of Pdyn-SPN clusters during all reward conditions. Each dot represents an individual tracked neuron that composes the cluster. (k) fraction of all tracked Pdyn-SPN neurons that belong to each functional cluster. (l) SPN clusters that were not consistently recruited during any of the reward conditions (averaged response to 20 trials of each reward condition). (m) SPN clusters that were consistently transiently inhibited during all reward conditions. (n) SPN clusters that were consistently transiently excited during the 0%, 3% and 10% conditions, but less for the 30% condition. (o) Penk-SPN cluster that was consistently transiently excited during 30% condition, moderately for the 0% condition and less for the 3% and 10% condition. (p) Pdyn-SPN cluster that was consistently transiently excited during the 3%, 10% and 30% sucrose conditions but less for the 0% (water) condition. (q) Pdyn-SPN cluster that was consistently transiently excited for all reward conditions. (r) Cartoon depicting multiclass linear Support Vector Classifier implementation (features = neurons, observations = mean activity during given trial, target = class of reward condition). (s) Mean of SVC coefficients for all Penk-SPNs (SVC features) of each functional cluster. (t) Mean of SVC coefficients for all Pdyn-SPNs (SVC features) of each functional cluster. (u) Linear SVC classification accuracy for every animal using mean SPN activity during reward epoch (0-8s from reward onset) and during control epoch (12-4s before reward onset) (n = 7 Penk-cre and 11 Pdyn-cre animals).

To measure the activity of mNAcSh SPNs during the consumption of variable reward conditions, we imaged the calcium dynamics of hundreds of individual SPNs using 2-photon fluorescence microscopy through endoscopic GRIN lenses. Preprodynorphin-cre (Pdyn-cre) and proenkephalin-cre (Penk-cre) animals were stereotaxically injected with 500 nl of AAV-DJ-Ef1α-DIO-GCaMP6s virus and implanted with a 600 μm diameter cylindrical GRIN lens in mNAcSh (**Fig. 1c-d**). After 4-6 weeks of recovery from surgery, animals were water restricted and trained to sip 10% sucrose reward at variable intervals while reversibly immobilized and head-restrained (**Fig. 1e**). All animals displayed consistent licking behavior while immobilized after 4 days of lick training and were then tested in 30 minute, 20 trial calcium imaging sessions each day for 4 consecutive days. In each session, animals consumed 1 of 4 possible sucrose reward conditions (0%, 3%, 10% or 30% sucrose in drinking water), with the order of condition days pseudorandomized and counterbalanced across animals (**Fig. 1f**). Animals immediately licked the presented sucrose reward, with an average licking bout duration of 7-8 seconds (**Fig. 1g**). There was no significant difference between reward conditions in the average amount of licking for each reward due to the small 10 ul reward volumes (**Fig. 1g**). This uniformity in lick behavior allowed for accurate comparison of SPN activity in response to the content of the different reward identities without any confound of differences in motor output (licking).

The resolution and low noise floor of 2-photon calcium imaging provides visualization of individual cell morphology and enabled high fidelity tracking of 715 individual SPNs across multiple days and behavioral conditions (**Fig. 2i-j, Extended Data Fig. 2a**). Individually tracked Pdyn-SPNs and Penk-SPNs displayed reward responses that were consistent for a given sucrose concentration, but that varied between the different sucrose concentrations (**Extended Data Fig. 2b-c**). Groups of individual SPNs were also correlated in their selectivity for specific sucrose concentrations. In order to group individual neurons together that had similar response profiles across the 4 sucrose conditions, k-means clustering was performed across all tracked SPNs. This unbiased approach to grouping neurons on the basis of similar activity revealed that there were 4 distinct clusters of Penk-SPNs and 5 distinct clusters of Pdyn-SPNs (**Fig. 1k-p, Extended Data Fig. 2d-g**). Notably, every cluster was composed of neurons from nearly all, if not all, of the mice that were tested, demonstrating that these functionally-defined mNAcSh clusters are generalizable across animals (**Fig. 2i-j, Extended Data Fig. 2b-c**). 32% of tracked Penk-SPNs and 41% of tracked Pdyn-SPNs were not consistently modulated by any of the reward conditions (**Fig. 1k**). A greater proportion of Penk-SPNs (32%) were inhibited in response to all 4 sucrose conditions than for Pdyn-SPNs (18%) (**Fig. 1l**). A large cluster of Penk-SPNs and Pdyn-SPNs were found to be active in response to the 0%, 3% and 10% sucrose conditions, but were relatively quiescent in response to the 30% sucrose condition (**Fig. 1m**). Nearly half of all reward-excited Penk-SPNs were selectively active far more for the high 30% sucrose condition (**Fig. 1n**). The presence of 3 robust and complimentary Penk-SPN clusters that are differently activated at the 30% condition indicates that there is a dramatic shift of mNAcSh Penk-SPN recruitment during high concentration reward that mice consume at a lower level relative to lower concentrations (**Fig. 1q-t**).

**Fig. 2.**
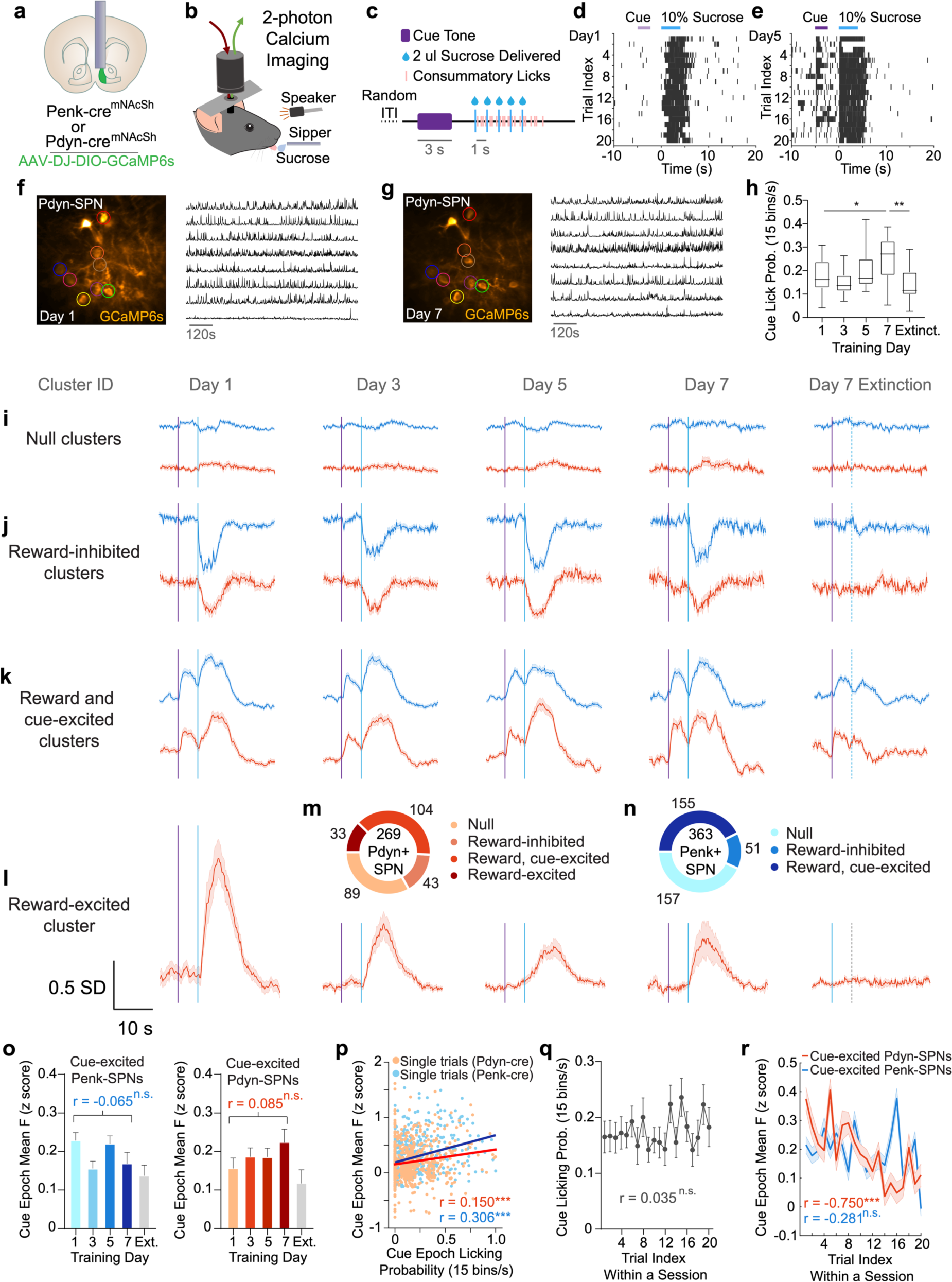
Reward-excited mNAcSh SPNS are cue-excited, but do not reflect reward expectation. (a) Coronal section cartoon displaying target brain region, cell-types and lens implant location. (b) Schematic of 2-photon calcium imaging through endoscopic lens during cue-reward Pavlovian conditioning. (c) Time-course of auditory cue and subsequent sucrose reward delivery for a given trial. (d and e) Representative lick raster depicting individual licks that were detected by contact licometer during cue and reward when animals were cue-naïve (d) and after conditioning (e). (f and g) Fluorescence time series of individual SPNs that were tracked from Pavlovian (f) day 1 session through (g) day 7 session. (h) Mean lick probability during cue epoch (0-5 seconds from cue-onset) for each day of Pavlovian conditioning and extinction (n = 16 animals; One-way ANOVA: Day 1 vs Day 7, *p = 0.020, Day 7 vs Extinction, **p = 0.002). (i) SPN clusters that were not consistently recruited during cue or reward throughout 7 days of Pavlovian conditioning (averaged response to 20 trials of each reward condition). (j) SPN clusters that were consistently transiently inhibited during reward throughout 7 days of Pavlovian conditioning. (k) SPN clusters that were consistently transiently excited during cue and reward throughout 7 days of Pavlovian conditioning and extinction. (l) Pdyn-SPN cluster that was consistently transiently excited during reward but not cue throughout 7 days of Pavlovian conditioning. (m) Fraction of all tracked Pdyn-SPN neurons that belong to each Pavlovian functional cluster. (n) Fraction of all tracked Penk-SPN neurons that belong to each Pavlovian functional cluster. (o) Mean cue response (0-5 seconds from cue-onset) of all cue-excited SPNs during each day of Pavlovian conditioning and extinction (n = 7 Penk-cre animals, 155 neurons; Pearson’s correlation: p = 0.105; n = 9 Pdyn-cre animals, 104 neurons; Pearson’s correlation: p = 0.083). (p) Correlation between mean cue response of all cue-excited cluster SPNs and mean cue epoch lick probability for every Pavlovian conditioning trial. Blue and red lines capture linear fit between cue response and cue licking for each cell-type. (Pearson’s correlation: ***p < 0.001). (q) Mean lick probability during cue epoch (0-5 seconds from cue-onset) for each trial within a session (Days 1, 3, 5 combined) (n = 16 animals; Pearson’s correlation: p = 0.288). (r) Mean cue response of all cue-excited cluster SPNs during each trial within a session (Days 1, 3, 5 combined) (n = 7 Penk-cre animals; Pearson’s correlation: p = 0.254; n = 9 Pdyn-cre animals; Pearson’s correlation: ***p < 0.001).

A distinct cluster of Pdyn-SPNs was also found to be active to all sucrose reward conditions (3%, 10% and 30%) but much less active to water (0% sucrose) (**Fig. 1o**). A small minority of Pdyn-SPNs (7%) were also found to be highly excited by every reward condition (**Fig. 1p**). To further quantify the specificity of mNAcSh SPN activity to the 4 reward preference conditions, a linear Support Vector Classifier (SVC) model was created. This SVC model was trained to predict which of the 4 sucrose conditions an animal consumed on a given trial only by considering the mean activity from each SPN during the reward epoch (**Fig. 1u**). The SVC performed with high accuracy and specificity for all 4 reward conditions for both Pdyn-SPNs and Penk-SPNs (**Fig. 1v-x**). The SVC model coefficients were greater for SPNs in the low and high concentration clusters for discriminating between 30% sucrose and the other reward conditions (**Fig. 1v-w**). This reaffirms that the primary differences in recruitment of reward-excited SPNs were occurring between the low and high reward concentration conditions. Altogether, these mNAcSh calcium imaging experiments across multiple reward preference conditions revealed substantial heterogeneity in the reward-evoked activity of individual SPNs. There are distinct mNAcSh SPN populations that are selective for preferred reward conditions and populations that are selective for unpreferred reward conditions. Reward-excited Penk-SPNs were highly selective in their reward response on the basis of reward concentration, while reward-excited Pdyn-SPNs were less selective for particular reward concentration conditions. Consistent recruitment of distinct mNAcSh SPN subpopulations to rewards of different preference suggests that these neurons may encode specific reward preference information.

### mNAcSh SPNs encode novelty of cues more than reward expectation

Since we observed heterogeneity in the activity of mNAcSh SPNs to rewards of different preference, we then sought to characterize potential differences in how individual mNAcSh SPNs respond to reward-predictive cues as a function of cue-reward associative learning. The same animals that were tested in the variable sucrose 2-photon imaging paradigm (see **Fig. 1**) were also trained to associate an auditory cue to subsequent delivery of 10% sucrose reward across 7 days of Pavlovian conditioning (**Fig. 2a-c**). Animals again displayed immediate licking in response to sucrose reward presentation, but initially did not show robust anticipatory licking to the cue (**Fig. 2d,h**). Animals began to display consistent anticipatory licking after multiple sessions of Pavlovian conditioning, indicating that the cue-reward association had been learned (**Fig. 2e,h**).

2-photon calcium imaging of mNAcSh SPNs expressing GCaMP6s occurred simultaneously to Pavlovian conditioning on training days 1, 3, 5 and 7, and 632 individual SPNs were successfully tracked across all Pavlovian imaging sessions (**Fig. 2f-g, Extended Data Fig. 3a**). K-means clustering was performed on 632 SPNs to stratify groups of neurons on the basis of similar cue and reward responses. This approach revealed that there were 3 distinct clusters of Penk-SPNs and 4 distinct clusters of Pdyn-SPNs within the context of the Pavlovian conditioning paradigm (**Fig. 2m-n, Extended Data Fig. 3b-g**). Not all reward-responsive mNAcSh SPNs (61% of all SPNs) were found to also be cue-responsive (41% of all SPNs) (**Fig. 2m-n**). There were also no cue-inhibited SPNs observed and all reward-inhibited SPNs had no consistent cue-response (**Fig. 2j**). There was also a distinct cluster of Pdyn-SPNs that was reward-excited but not cue-excited (**Fig. 2l**), while all reward-excited Penk-SPNs were also cue-excited (**Fig. 2k**). The Pdyn-SPN cluster that was only reward-excited also showed smaller reward-excitation to the 10% sucrose reward in later Pavlovian conditioning days, suggesting that the expectation of reward diminished the magnitude of reward-excitation (**Fig. 2l**). To test for the persistence of cue and reward-responses during the absence of expected reward delivery, the latter half (last 10 out of 20 trials) of trials on Pavlovian day 7 were reward omission trials, where the cue is presented as usual but the expected sucrose reward is not delivered. The SPNs that were found to be exclusively reward-responsive but not cue-responsive showed no response during Pavlovian extinction trials when reward was omitted (**Fig. 2j,l**), further suggesting that the reward-responses were truly specific to the content of the reward itself. However, cue and reward-excited SPNs displayed a diminished, but persistent cue and reward-excitation during the 10 extinction trials (**Fig. 2k**).

Surprisingly, the magnitude of cue-excitation of the cue-excited SPN clusters did not increase across Pavlovian training days (**Fig. 2o**) as is canonically observed in VTA DA neurons during cue-reward associative learning^27,28^, or as was recently observed via one-photon calcium imaging in SPNs expressing the dopamine D2 receptor^29^. We also found that the magnitude of cue-excitation was only weakly correlated with amount of anticipatory licking per trial (**Fig. 2p**). Taken together, these results demonstrate that mNAcSh SPN cue-responses do not robustly change as a function of cue-reward associative learning and do not reflect reward expectation. Additionally, Pdyn-SPNs decreased the magnitude of their cue-excitation across trials within an imaging session, suggesting that the cue-response of Pdyn-SPNs may reflect the salience of the cue tone rather than its motivational value (**Fig. 2q-r**).

### Stability and overlap of sucrose concentration and pavlovian neuronal ensemble clusters

Given reports that striatal SPNs display stochastic encoding at a trial-by-trial level of valence (Domingues et al. 2023 co-submission), we sought to evaluate whether the distinct SPN ensembles we identified, whose means robustly predicted sucrose concentration trial type (**Fig. 1r-u**), displayed similar or different activity patterns between trials and across paradigms. This may represent a unified encoding strategy for striatal D1- and D2-SPNs. We reliably tracked hundreds of Penk-SPNs during 4 variable sucrose concentration sessions (0%, 3%, 10%, and 30%) as well as 5 pavlovian conditioning sessions (Day 1, 3, 5, 7 and extinction). Across both paradigms, cell responsivity was largely consistent. Penk-SPNs that were identified as having increased activity in response to some or multiple sucrose concentrations were also largely activated in response to both the cue and sucrose reward during pavlovian conditioning (**Fig. 3a-b**). Most variable sucrose inhibited neurons were also inhibited during pavlovian conditioning (**Fig. 3a-b**). Pdyn-SPNs displayed similar trends overall, with the various sucrose activated clusters (sucrose low, sucrose, and all rewards) displaying activation in response to the cue and reward during pavlovian conditioning (**Fig. 3c-d**). An additional pavlovian reward excited-only was identified for Pdyn-SPNs, and the majority of these neurons were spread across variable sucrose-activated clusters, though interestingly some overlapped with the null and even inhibited variable sucrose clusters (**Fig. 3d**). Overall, the sign (activated or inhibited) of neural responses for both Penk- and Pdyn-SPNs across clusters was consistent across both behavioral paradigms.

**Fig. 3.**
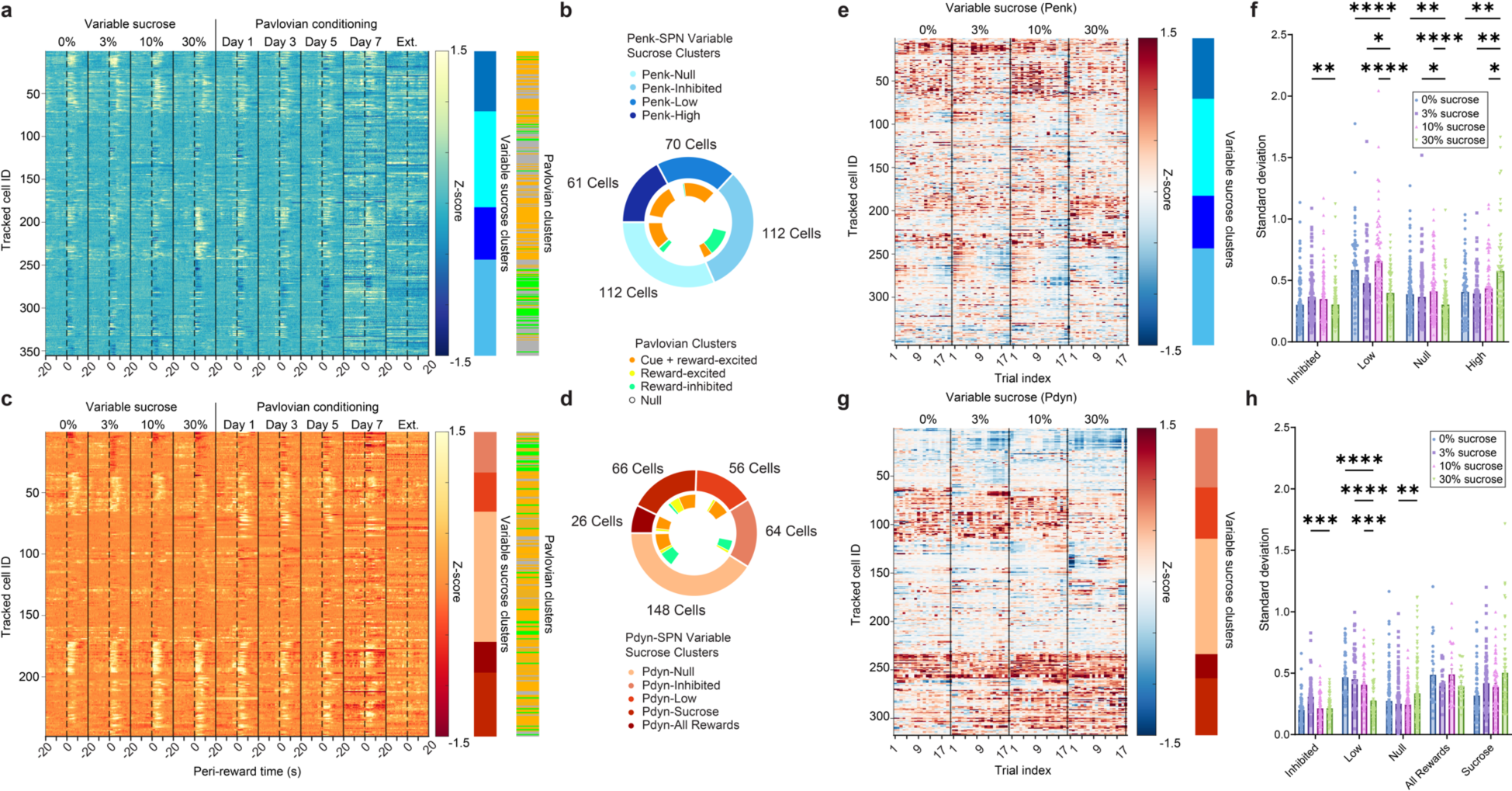
Stability and overlap of Penk- and Pdyn-SPN reward and pavlovian clusters. (a) Penk-SPNs tracked across each variable sucrose (0%, 3%, 10%, 30%) and pavlovian conditioning (Day 1, 3, 5, 7 and Extinction) session. Each row represents a tracked neuron activity aligned to reward delivery across paradigms (dotted line). Heatmaps represent cluster identity for variable sucrose and pavlovian conditioning experiments. (b) Fraction of Penk-SPNs in each variable sucrose cluster (outer donut) that were found to be cue or reward-responsive (inner donut) within the context of the Pavlovian conditioning paradigm. (c) Penk-SPNs tracked across each variable sucrose (0%, 3%, 10%, 30%) and pavlovian conditioning (Day 1, 3, 5, 7 and Extinction) session. Each row represents a tracked neuron activity aligned to reward delivery across paradigms (dotted line). Heatmaps represent cluster identity for variable sucrose and pavlovian conditioning experiments. (d) Mean trial activity of Penk-SPNs during the consumption period (0 to 2s post delivery) for each trial in each session (20 delivery trials per session). Cells are matched across sucrose concentration and sorted by cluster identity. (f) Comparison of standard deviation for each neuron across trials for each sucrose concentration as a measure of variability in response to stimulus presentation (Two-way repeated measures ANOVA, main effect of cluster *p*<0.0001, main effect of concentration *p*<0.0001, interaction between cluster and concentration *p*<0.0001; Asterisks indicate significant Bonferroni’s post-hoc test, \*\*\*\**p*<0.0001, \*\*\**p*<0.001, \*\**p*<0.01, \**p*<0.05). (g) Mean trial activity of Pdyn-SPNs during the consumption period (0 to 2s post delivery) for each trial in each session (20 delivery trials per session). Cells are matched across sucrose concentration and sorted by cluster identity. (h) Comparison of standard deviation for each neuron across trials for each sucrose concentration as a measure of variability in response to stimulus presentation (Two-way repeated measures ANOVA, main effect of cluster *p*<0.0001, no main effect of concentration *p*=0.48, interaction between cluster and concentration *p*<0.0001; Asterisks indicate significant Bonferroni’s post-hoc test, \*\*\*\**p*<0.0001, \*\*\**p*<0.001, \*\**p*<0.01, \**p*<0.05).

We next asked whether there were any differences in variability in response across trials in variable sucrose response for Penk- and Pdyn-SPNs. We calculated the mean fluorescence response within the reward window (0 to 2s) for each trial in each session (**Fig. 3e,g**). We then calculated the standard deviation (σ) across trials within a single concentration as a measure of variability in neural response (**Fig. 3f**). For Penk-SPNs, we find that responses to the preferred concentrations (0%, 3%, 10%) of sucrose to be minimally variable and stable across concentrations (**Fig. 3f**). By contrast, the stability of the response to 30% sucrose was significantly different relative to other sucrose concentrations for several clusters. For the sucrose low and sucrose null clusters, the response to 30% sucrose was significantly less variable than the other concentrations. Interestingly, for the high sucrose concentration cluster, which displayed elevated activity in response to the non-preferred 30% sucrose, variability across trials was elevated relative to other sucrose concentrations (**Fig. 3f**). This supports the contention that Penk-SPNs may preferentially encode the non-preferred high sucrose concentration. Pdyn-SPN activity was also quite consistent across concentrations within clusters, and overall displayed less differences relative to sucrose concentration compared to the Penk-SPNs (**Fig. 3h**). Only the Pdyn-SPN low sucrose concentration cluster displayed similar concentration dependent changes in variability, with 30% sucrose producing the least variable response across trials. This result suggests that both Penk- and Pdyn-SPNs play a role in encoding the rewarding properties of lower, more palatable concentrations of sucrose.

### Functionally defined mNAcSh SPN clusters are anatomically separated along anterior-posterior axis

Previous studies have demonstrated stark differences in the molecular identity and function of SPNs, even within the mNAcSh ^2,11^. Accordingly, the functionally defined SPN clusters that were identified in the variable sucrose and Pavlovian conditioning behavioral paradigms (see **Fig. 1-2**) were analyzed to relate the functional identity of SPNs to their relative position within the focal plane during in vivo 2-photon calcium imaging. The x and y position of each tracked SPN in the focal plane was calculated using the ROIs of the identified neuron. The distribution of x and y positions in the focal plane for each cluster was considered and mapped to the anterior-posterior and medial-lateral orientation of the animal during the imaging session. The implanted endoscopic lenses were 600 um in diameter and thus, segregation of functional SPN clusters within the imaging focal plane represents hundreds of microns of anatomical separation within mNAcSh. Visualizing the spatial distribution of functionally defined SPN clusters revealed that reward-inhibited SPNs were specific to anterior portions of the focal plane, for the variable sucrose and Pavlovian imaging sessions (**Fig. 4a-j**). This strong divergence in reward response of mNAcSh SPNs across the anterior-posterior axis may indicate a critical difference in the function of these anatomically separable SPN populations.

**Fig. 4.**
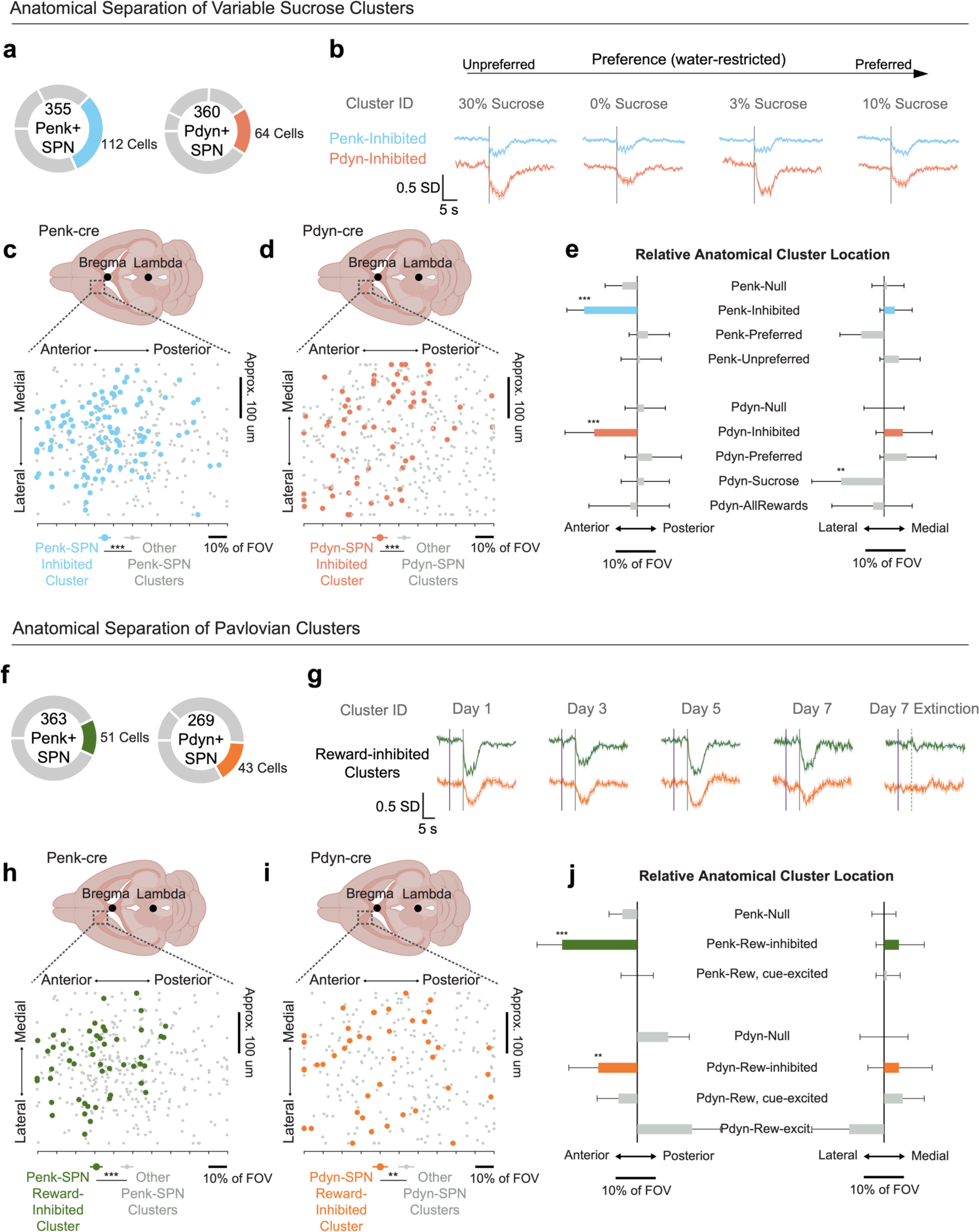
Reward-inhibited SPN clusters are anatomically anterior to other SPN clusters. (a) Fraction of all tracked Penk-SPNs and Pdyn-SPNs that are part of reward-inhibited clusters. (b) Mean activity of SPN clusters that were consistently transiently inhibited during all variable sucrose reward conditions. (c) Position of individual Penk-SPNs in focal plane during in vivo 2-photon calcium imaging sessions; arranged by variable sucrose response cluster (n = 4 clusters, 7 animals; one-way ANOVA: ***p < 0.001). (d) Position of individual Pdyn-SPNs in focal plane during in vivo 2-photon calcium imaging sessions; arranged by variable sucrose response cluster (n = 5 clusters, 11 animals; one-way ANOVA: ***p < 0.001). (e) Relative anatomical location of SPN variable sucrose response clusters. (f) Fraction of all tracked Penk-SPNs and Pdyn-SPNs that are part of Pavlovian reward-inhibited clusters. (g) Mean activity of SPN clusters that were consistently transiently inhibited during reward throughout 7 days of Pavlovian conditioning. (h) Position of individual Penk-SPNs in focal plane during in vivo 2-photon calcium imaging sessions; arranged by Pavlovian conditioning response cluster (n = 3 clusters, 7 animals; one-way ANOVA: ***p < 0.001). (i) Position of individual Pdyn-SPNs in focal plane during in vivo 2-photon calcium imaging sessions; arranged by Pavlovian conditioning response cluster (n = 4 clusters, 9 animals; one-way ANOVA: **p = 0.004). (j) Relative anatomical location of SPN Pavlovian conditioning response clusters.

### mNAcSh-VP^Penk^ parallels selectivity and cue-excitation of the Penk-SPN high concentration neuronal ensemble cluster

Our results thus far show greater heterogeneity in responses of Penk-SPNs to reward concentration, with a large cluster recruited specifically during consumption of the 30% reward condition which mice consume less readily than lower concentrations. In contrast, Pdyn-SPNs were less heterogenous to reward preference (**Fig. 1 and Extended Fig. 1**). Prior studies have identified and characterized two major output pathways from the mNAcSh: D2/Penk-SPNs projecting to ventral pallidum (VP), and D1/Pdyn-SPNs projecting to lateral hypothalamus (LH), and suggest that they are necessary for reward consumption^30^. To determine whether they play a role in relative reward preference we tested whether the stratification of reward preference information in mNAcSh is propagated selectively to these downstream brain regions via major mNAcSh efferent projections. We retrogradely traced the major output projections from mNAcSh Penk-SPNs and Pdyn-SPNs using retrobeads and fluorescent in situ hybridization (FISH) (**Extended Data Fig. 6a**,d). It was also found that Pdyn-SPNs densely project to LH and sparsely to VTA with minimal collateralization (7.3%) (**Extended Data Fig. 6a-c**), as has been previously reported of D1R expressing SPNs ^31^. In congruence with previous literature ^32^, Penk-SPNs were found to densely project to ventral pallidum (VP) and less densely to lateral hypothalamus (LH) and these circuits had minimal collateralization (13.3%) (**Extended Data Fig. 6d-f**).

To determine if these major SPN efferent circuits were selectively recruited to particular reward preference conditions and not others, we conducted fiber photometry recordings while animals consumed the 4 reward preference conditions (0%, 3%, 10% or 30% sucrose; **Fig. 5 and Extended Data Figure 7**). Penk-cre and Pdyn-cre mice were first injected with AAV-DJ-Ef1α-DIO-GCaMP6s in the mNAcSh and then implanted 400 um diameter fiber optic cannulas into the VP and LH (Penk-cre) or LH and VTA (Pdyn-cre) (**Fig. 5a and Extended Data Fig. 6**). We expect that canonical projections (e.g. NAcSh-VP^Penk^ and mNAcSh-LH^Pdyn^; **Fig. 5e-h, i-l**), which we verified (**Extended Data Fig. 6**) would be consistent with reward and cue responsivity observed at the single cell level (**Fig.1 & 2**) while less dense and established projections (e.g. NAcSh-LH^Penk^ and mNAcSh-VTA^Pdyn^, **Extended data Fig. 7a-h, i-p**) would not. To ensure uniformity in licking behavior identical to the 2-photon calcium imaging experiments, photometry animals were tested in the same head-restrained behavior paradigm (see **Fig. 1**) (**Fig. 5a-b**).

**Fig. 5.**
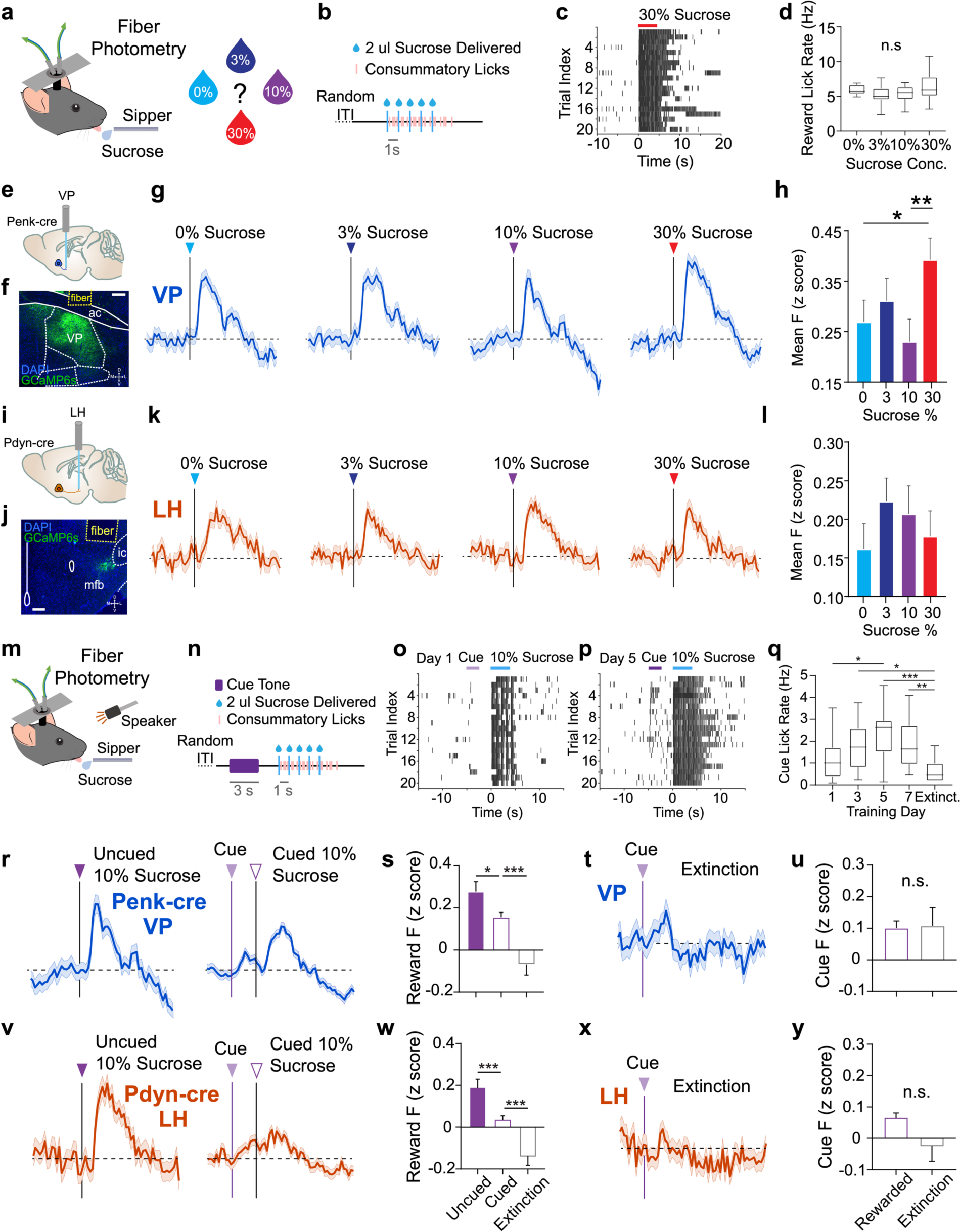
mNAcSh-VP^Penk^ parallels unpreferred reward selectivity and cue-excitation of Penk-SPN cluster. (a) Schematic of simultaneous fiber photometry during consumption of variable sucrose reward. (b) Time-course of sucrose reward delivery for a given trial. Droplets were either 0%, 3%, 10% or 30% sucrose in water. (c) Representative lick raster depicting individual licks that were detected by contact licometer during behavior and photometry session. (d) Mean lick rate during reward epoch (0-8 seconds from reward onset) for each sucrose reward condition (n = 15 animals; one-way ANOVA: p = 0.280). (e) Sagittal section cartoon displaying target injection region, cell-type and fiber implant locations. (f) Representative coronal section showing Penk-SPN terminals in VP expressing GCaMP6s underneath the fiber implant (scale bar = 200 um). (g) Mean mNAcSh-VP^Penk^ reward-response for every trial of each variable sucrose reward condition. (h) Mean mNAcSh-VP^Penk^ activity over reward response epoch (2-12 seconds from reward-onset) for each condition (n = 9 animals, 190+ trials; one-way ANOVA: 30% vs 0%, *p = 0.044, 30% vs 10%, **p = 0.009). (i) Sagittal section cartoon displaying target injection region, cell-types and fiber implant locations for mNAcSh-LH^Pdyn^ recordings. (j) Representative coronal section showing Pdyn-SPN terminals in LH expressing GCaMP6s underneath the fiber implant (scale bar = 150 um). (k) Mean mNAcSh-LH^Pdyn^ reward-response for every trial of each variable sucrose condition. (l) Mean mNAcSh-LH^Pdyn^ activity over reward response epoch (2-12 seconds from reward-onset) for each condition (n = 6 animals, 123+ trials, one-way ANOVA: p = 0.550). (m) Schematic of simultaneous fiber photometry during Pavlovian cue-reward conditioning. (n) Time-course of auditory cue and subsequent sucrose reward delivery for a given trial. (o and p) Representative lick raster depicting individual licks that were detected by contact licometer during cue and reward when animals were cue-naïve (M) and after conditioning (N). (q) Mean lick rate during cue epoch (0-5 seconds from cue-onset) for each day of Pavlovian conditioning and extinction (n = 15 animals; one-way ANOVA: Day 1 vs Day 5, *p = 0.041, Day 3 vs Ext., *p = 0.011, Day 5 vs Ext., ***p < 0.001, Day 7 vs Ext., **p = 0.005). (r) Mean mNAcSh-VP^Penk^ reward-response for every uncued 10% sucrose trial and every cued 10% sucrose trial. (s) Mean mNAcSh-VP^Penk^ activity over reward response epoch (2-12 seconds from reward-onset) for uncued 10% sucrose, cued 10% sucrose and extinction trials (n = 9 animals, 96+ trials; one-way ANOVA: Uncued vs Cued, *p = 0.014, Cued vs Ext., ***p < 0.001). (t) Mean mNAcSh-VP^Penk^ reward-response for every trial of day 7 Pavlovian extinction. (u) Mean mNAcSh-VP^Penk^ activity over cue response epoch (2-5 seconds from cue-onset) for cued 10% sucrose and extinction trials (n = 9 animals, 96+ trials; two sample t-test: p = 0.909). (v) Mean mNAcSh-LH^Pdyn^ reward-response for every uncued 10% sucrose trial and every cued 10% sucrose trial. (w) Mean mNAcSh-LH^Pdyn^ activity over reward response epoch (2-12 seconds from reward-onset) for uncued 10% sucrose, cued 10% sucrose and extinction trials (n = 6 animals, 63+ trials, one-way ANOVA: Uncued vs Cued, ***p < 0.001, Cued vs Ext., ***p < 0.001). (x) Mean mNAcSh-LH^Pdyn^ reward-response for every trial of day 7 Pavlovian extinction. (y) Mean mNAcSh-LH^Pdyn^ activity over cue response epoch (2-5 seconds from cue-onset) for cued 10% sucrose and extinction trials (n = 6 animals, 63+ trials, two sample t-test: p = 0.073).

mNAcSh-VP^Penk^ axon terminals had reward-excitation to all 4 sucrose preference conditions. However, the magnitude of mNAcSh-VP^Penk^ reward-excitation was significantly greater for the higher concentration reward condition than the lower reward conditions (**Fig. 5g-h**). This greater excitation to the unpreferred condition directly parallels the preference selectivity of the Penk-High (30% sucrose) cluster we identified during 2-photon imaging experiments (see **Fig. 1n**). By contrast, no significant difference in activation as a function of reward concentration was observed in the less dense mNAcSh-LH^Penk^ pathway (**Extended data Fig. 7c-d**), suggesting that the large high-reward activated Penk-SPN cluster may preferentially project to the VP and not the LH. Similar to what we observed at the single cell level in the mNAcSh (**Fig. 1p**), the mNAcSh-LH^Pdyn^ projection was not selective for any particular reward concentration (**Fig. 5k-l**), with equal increases in activity observed across concentrations. Overall, responses of mNAcSh-VTA^Pdyn^ projections, which we found to be relatively sparse (**Extended Data Fig. 6c**), were minimal across sucrose concentrations (**Extended Data Fig. 7k-l**), though increased fluorescence was observed at 3% sucrose relative to 10% sucrose (**Extended Data Fig. 7l**). Importantly, no significant activation of either major projection (NAcSh-VP^Penk^ and mNAcSh-LH^Pdyn^) was observed during bouts of locomotion (**Extended Data** Figure 9a-g). Reward preference selectivity of some mNAcSh efferents and not others suggests that the heterogeneous reward preference encoding observed in mNAcSh SPN populations is selectively propagated downstream through particular efferent structures.

To examine whether this pathway specificity persisted during conditioning, these animals were also tested in a 7 day Pavlovian conditioning paradigm identical to the head-restrained paradigm used during 2-photon calcium imaging (see **Fig. 2b-c**). Activity of both mNAcSh-VP^Penk^ and mNAcSh-LH^Pdyn^ projections were found to be cue-excited throughout Pavlovian conditioning in addition to having consistent 10% sucrose reward-excitation, though the cue response was significantly reduced compared to the uncued response to 10% sucrose for both projections (**Fig. 5r-u, v-y**). Furthermore, cue-excitation of mNAcSh-VP^Penk^ persisted even during the 10 extinction trials where the reward was omitted (**Fig. 5t-u**) while the excitation of mNAcSh-LH^Pdyn^ projections did not persist during extinction (**Fig. 5x-y**). This mNAcSh-VP^Penk^ response profile within the Pavlovian conditioning paradigm parallels the cue and reward-excitation that was observed in Penk-SPNs cue and reward-excited clusters (see **Fig. 2k, Extended Data Fig. 5**). Given our calcium imaging-based characterization of Penk-SPNs in mNAcSh, this suggests that the mNAcSh-VP^Penk^ circuit is likely largely composed of the unpreferred reward selective Penk-SPN subpopulation in mNAcSh. Thus, the separation of distinct reward preference information streams in mNAcSh is continued through the mNAcSh-VP^Penk^ circuit, which selectively encodes unpreferred rewards.

### mNAcSh-VP^Penk^ causally drives low positive reward value

mNAcSh-VP^Penk^ was found to have greater reward-excitation to the 30% sucrose concentration reward condition than the lower concentration conditions, reflecting what we found at the single neuron level in a large and distinct cluster of mNAcSh Penk-SPNs. However, it remained unclear whether mNAcSh-VP^Penk^ reward-related activity causally contributes to consummatory behavior for high concentration, unpreferred rewards. To test the role of mNAcSh-VP^Penk^ in casually driving preference for reward, we injected Penk-cre mice with 300 nl of AAV5-EF1α-DIO-parapinopsin(PPO)-Venus ^33^ or AAV5-EF1α-DIO-ChR2-eYFP bilaterally into mNAcSh and implanted a 200 um diameter fiber optic bilaterally into VP (**Fig. 6e-f,i-j**). After recovering from surgery, animals were water-restricted and trained to lick 10% sucrose reward while reversibly immobilized and head-restrained (**Fig. 6a**). After 4 days of lick training, all animals displayed consistent lick behavior in response to variable interval sucrose presentation and were then put through 2 consecutive test days.

**Fig. 6.**
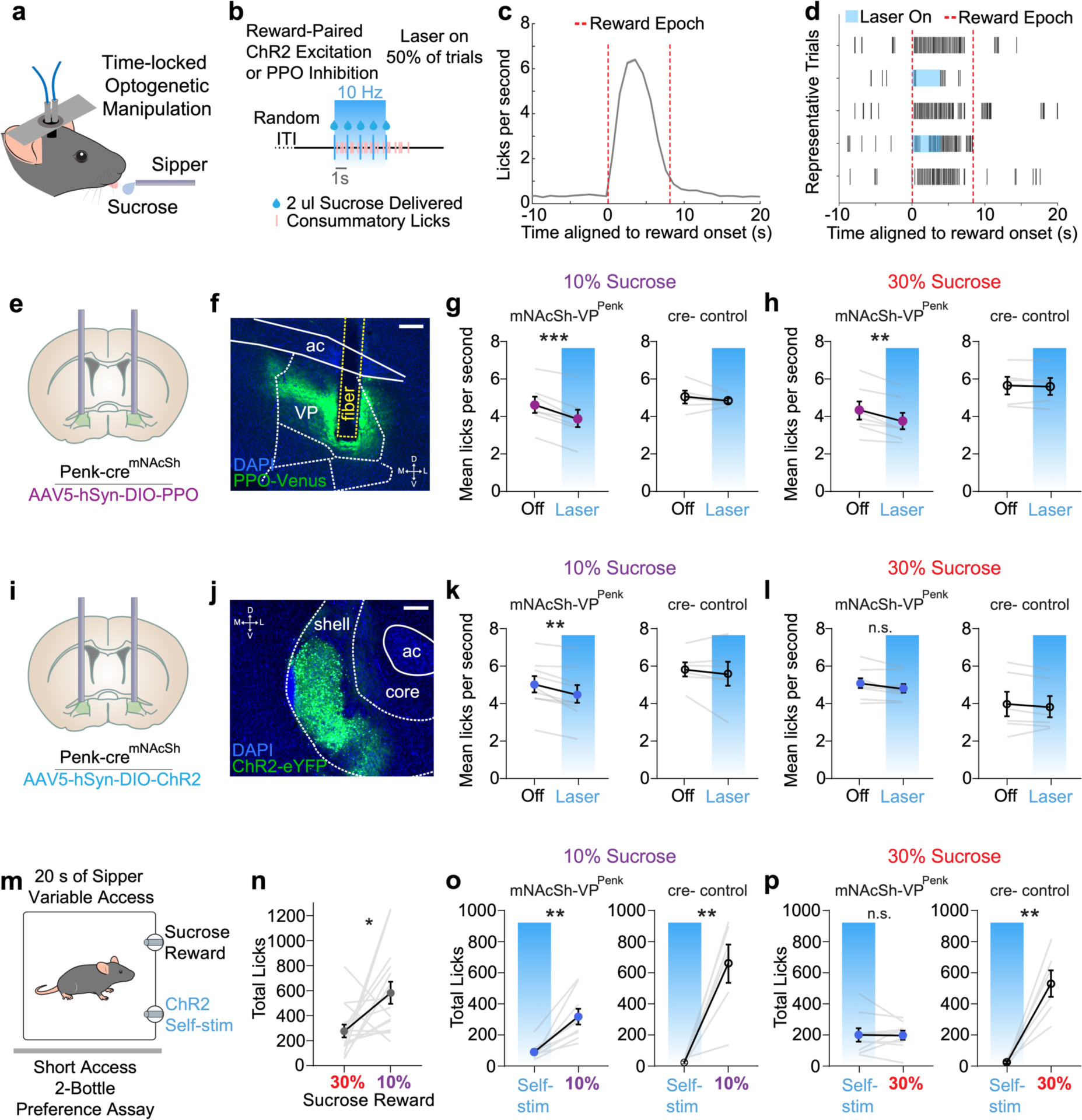
mNAcSh-VP^Penk^ causally drives low-positive reward preference. (a) Schematic of timelocked-optogenetic neural activity manipulation during consumption of variable sucrose reward. (b) Time-course of sucrose reward delivery for a given trial. 50% of trials also had 4 seconds of 10 Hz optogenetic stimulation concurrent with reward delivery. (c) Lick rate distribution relative to reward-onset for all conditions. Comparison of lick rate between conditions was restricted to the indicated reward epoch (0-8 seconds from reward-onset). (d) Representative lick raster displaying individual measured licks during trials without stimulation and trials with stimulation. (e) Coronal section cartoon displaying target injection region, cell-type and bilateral fiber implant location. (f) Representative coronal section showing Penk-SPN terminals in VP expressing PPO underneath optic fiber implant (scale bar = 200 um). (g) Within-session comparison of 10% sucrose consumption between trials with and without reward-paired optogenetic inhibition (n = 7 treatment, 5 control animals; pairwise t-test: ***p < 0.001, p = 0.418). (h) Within-session comparison of 30% sucrose consumption between trials with and without reward-paired optogenetic inhibition (n = 7 treatment, 5 control animals; pairwise t-test: **p = 0.003, p = 0.619). (i) Coronal section cartoon displaying target injection region, cell-type and bilateral fiber implant location. (j) Representative coronal section showing Penk-SPN terminals in VP expressing ChR2 underneath optic fiber implant (scale bar = 200 um). (k) Within-session comparison of 10% sucrose consumption between trials with and without reward-paired optogenetic excitation (n = 9 treatment, 6 control animals; pairwise t-test: **p = 0.003, p = 0.437). (l) Within-session comparison of 30% sucrose consumption between trials with and without reward-paired optogenetic excitation (n = 9 treatment, 6 control animals; pairwise t-test: p = 0.057, p = 0.327). (m) Preference between 10% sucrose reward sipper and 30% sucrose reward sipper (n = 15 animals; pairwise t-test: *p = 0.021). (n) Schematic of brief access 2-bottle choice task: animal makes choice between sucrose reward sipper and lick-triggered 2 second, 10 Hz optogenetic self-stimulation. (o) Preference between 10% sucrose reward and optogenetic self-stimulation (n = 9 treatment, 6 control animals; pairwise t-test: **p = 0.001, **p = 0.004). (p) Preference between 30% sucrose reward and optogenetic self-stimulation (n = 9 treatment, 6 control animals; pairwise t-test: p = 0.956, **p = 0.002).

Each test day involved one 30 minute, 20 trial session in which the animals were presented with either 10% or 30% sucrose reward. The order of sucrose condition days was pseudorandomized and counterbalanced across animals. For consistency with the imaging and photometry behavior paradigms, animals were presented with 5 droplets of sucrose reward per trial with a total aggregate volume of 10 ul and 1 second between droplet delivery (**Fig. 6b**). Animals immediately licked the presented sucrose reward, with an average licking bout duration of 7-8 seconds (**Fig. 6c**). On a pseudorandom 50% of the trials within a session, time-locked optogenetic manipulation of mNAcSh-VP^Penk^ activity would be paired with reward delivery, via 4 seconds of 10 Hz blue light delivery (**Fig. 6b**). 10Hz optogenetic stimulation frequency was chosen because it has been reported previously to be a physiologically relevant level of neural activity for SPNs during phasic reward-related activity ^12,13^. Any differences in consummatory licking between no laser and laser trials were recorded via a contact lickometer with millisecond precision (**Fig. 6d**) and all differences were evaluated within subject to account for the baseline variability in lick rate between animals. To control for possible behavioral effects from the delivery of the blue light itself, cre-littermate controls were also injected, implanted, and tested in identical conditions.

Time-locked photoinhibition (PPO-injected mice) of mNAcSh-VP^Penk^ during consumption significantly reduced the reward epoch lick rate regardless of whether animals consumed 10% or 30% sucrose reward (**Fig. 6g-h**). This demonstrates that mNAcSh-VP^Penk^ reward-excitation is required for normal consummatory licking, in accordance with previous findings (Carina and Ana, 2022). Since mNAcSh Penk-SPNs specifically encode reward preference information and mNAcSh-VP^Penk^ is reward-excited, the reduction in consummatory licking during mNAcSh-VP^Penk^ photoinhibition is likely due to complete disruption of reward information in this circuit.

Time-locked photoexcitation of mNAcSh-VP^Penk^ during consumption impacted consummatory licking that was reward-dependent. Time-locked photoexcitation of mNAcSh-VP^Penk^ reduced the consummatory licking of preferred 10% sucrose reward (**Fig. 6k**) but did not change the consumption of unpreferred 30% sucrose reward (**Fig. 6l**). This lack of effect is likely due to the notion that mNAcSh-VP^Penk^ already has higher endogenous activity during the consumption of 30% sucrose reward (see **Fig. 5**) thus suggest exogenous photoexcitation is not capable of elevating this activity above this ceiling to further to alter behavior. By contrast, in the preferred reward condition, mNAcSh-VP^Penk^ has less endogenous activity and 10 Hz exogenous photoexcitation would more profoundly changes the activity of the circuit.

Notably, mNAcSh-VP^Penk^ exogenous photoexcitation is not disrupting the motor action of licking itself, as no disruption of licking behavior was observed during the consumption of 30% sucrose reward (**Fig. 6l**). To isolate any possible effects from motor action impairment, a set of additional experiments were designed in which mNAcSh-VP^Penk^ was photoexcited or photoinhibited as animals freely moved through a behavioral arena in red light conditions (**Extended Data Fig. 9h-j**). Movement velocity was not changed by mNAcSh-VP^Penk^ photoinhibition (**Extended Data Fig. 9k**), nor by mNAcSh-VP^Penk^ photoexcitation (**Extended Data Fig. 9m**), further suggesting that manipulation of this circuit during headfixed licking behavior was influencing the processing of reward information and not disrupting the motor execution of licking.

Interestingly, no effect of silencing mNAcSh-LH^Pdyn^ projections was observed for either 10% of 30% sucrose delivery (**Extended Data Fig. 9o-q**). However, a selective reduction in licking was observed when activation of the mNAcSh-LH^Pdyn^ projection was paired with 10% sucrose (**Extended Data Fig. 9r-s**) but not 30% sucrose (**Extended Data Fig. 9t**). Both at the single cell (**Fig. 1**) and projection level (**Fig. 5**), Pdyn neuron and projection activity was not consistently associated with selective reward concentrations, though graded responses existed. Recently, a study showed that Pdyn/D1-expressing NAc neurons enhanced LH glutamatergic activity^34^, which has previously been shown to be aversive^35^. Hence, it is possible that activating mNAcSh-LH^Pdyn^ projections could yield a reduction in licking observed in the 10% sucrose concentration condition. There was no significant effect of mNAcSh-LH^Pdyn^ inhibition or activation on locomotion (**Extended Data Fig. 9l**,n).

We next posited that if mNAcSh-VP^Penk^ is causally driving a low positive reward preference, then photoexcitation of this circuit alone should be inherently rewarding. To test this hypothesis, we tested mice in the 2-bottle choice paradigm (see **Extended Data Fig. 1**). mNAcSh-VP^Penk:ChR2^ expressing animals were trained to lick 10% sucrose reward from two sucrose spouts within a behavioral chamber. After training the mice to approach and lick the sucrose spouts for 5 days, we put the animals through 6 consecutive test days. Test days 1, 3, 5 were ‘reset’ days to neutralize any left versus right biases that resulted from other test conditions as described before (see **Extended Data Fig. 1**). On test day 2, animals were given brief access to both a 10% and a 30% sucrose sipper. In accordance with our earlier results (see **Extended Data Fig. 1**), mNAcSh-VP^Penk:ChR2^ animals showed a preference for 10% sucrose over 30% sucrose (**Fig. 6m**). On test days 4 and 6, the animals were again tested for their preference between two sippers. However, one of the two sippers contained no liquid and licking it triggered 2 seconds of 10 Hz closed loop optogenetic photoexcitation (**Fig. 6n**). This stimulation duration and frequency was chosen to reasonably recapitulate mNAcSh-VP^enk^ activity during 30% sucrose reward licking. The other sipper that the animals could choose instead of the self-stimulation sipper contained either 10% or 30% sucrose. The order of test days in which animals got the 10% or 30% alternative sipper and the sipper location were randomized and counterbalanced across animals. This experiment was designed to test whether animals find mNAcSh-VP^Penk^ photoexcitation inherently rewarding and how the reinforcement of mNAcSh-VP^Penk^ photoexcitation compares to the 10% and 30% sucrose consummatory conditions. We found that mNAcSh-VP^Penk:ChR2^ animals licked the empty self-stimulation sipper significantly more than cre-controls, that mNAcSh-VP^Penk^ photoexcitation is inherently reinforcing. Furthermore, cre+ animals licked the 10% sucrose sipper significantly more than the self-stimulation sipper (**Fig. 6o**) but showed no significant preference between the 30% sucrose and the self-stimulation sippers (**Fig. 6p**). This experiment indicates that the relative preference of exogenous mNAcSh-VP^Penk^ excitation is comparable to that of 30% sucrose reward, but measurably lower than the 10% sucrose reward. Overall, these complementary experiments demonstrate that mNAcSh-VP^Penk^ causally drives low positive reward preference. This leads us to conclude that not only is mNAcSh-VP^Penk^ endogenously recruited during the consumption of unpreferred rewards, but that endogenous activity during reward likely acts to influence the drive for perceived reward preference and consumption.

## Discussion

Here we report that distinct mNAcSh SPN ensembles encode relative reward preference. Separate subpopulations of mNAcSh SPNs are recruited during the consumption of preferred rewards and unpreferred rewards. We find that this stratification of reward preference information in mNAcSh is then propagated through efferent projections, with unpreferred reward signals relayed to VP by Penk-SPNs. Photo-activation of the mNAcSh-VP^Penk^ circuit was found to cause no change in consummatory behavior when the animals were consuming unpreferred reward but caused reductions in the consumption of preferred reward. Self-stimulation experiments further revealed that mNAcSh-VP^Penk^ activation itself was inherently rewarding and elicited consummatory behavior approximately equivalent to the lower preference natural reward. These findings present a mechanism for how mNAcSh likely integrate incoming reward-related information, represents reward information on the basis of relative preference and act coordinate the appropriately motivated consummatory behavior.

It remains unresolved how much separation of reward identity, their subsequent preference and the underlying relative value attributed to these rewards is present in brain regions upstream of accumbens and which afferents are supplying this information. There is recent evidence that neurons in VP encode reward value and project to NAc ^7,8^. Furthermore, reward value signals in VP may even be present earlier than in NAc ^6^. These previous findings and our current results suggest that some reward value information may be originally transmitted from VP to mNAcSh, where it is further amplified and propagated.

In addition to the evidence for separation of reward information upstream of mNAcSh, the stratification of this information could be further enhanced locally due to established interconnectivity between SPNs in mNAcSh^36^. Previous studies show a high degree of interconnectivity between nearby SPNs in NAc ^37,38^. The divergence in SPN recruitment on the basis of reward preference is likely further organized due to local inhibition of preferred reward SPNs by unpreferred reward SPNs when they are selectively activated.

The observation that reward-inhibited SPNs are anterior to other cue and reward-excited SPN clusters (see **Fig. 4**) is consistent with recent studies showing that incoming glutamatergic innervation from basolateral amygdala, ventral hippocampus and medial thalamus to anterior NAc is decreased during reward consumption and glutamatergic innervation to posterior NAc is increased during reward consumption ^39^. This finding strongly suggests that differences in glutamatergic innervation in NAc along the anterior-posterior axis result in differences in SPN reward processing and that the anterior-posterior division is a continuum of individual neurons rather than a strict boundary. Divergence in the reward-related activity of NAc populations has also previously been observed across the mediolateral and dorsoventral axes ^11,40^.

Reward and cue-related signaling in NAc core has been found to be in part mediated by mesolimbic dopamine release ^17,41^. However, it remains unclear how the differences in dopaminergic innervation of NAc core and mNAcSh contribute to differences in reward processing. Our current results demonstrate that many Penk-SPNs, that largely co-express inhibitory D2Rs ^2,42^, are excited during reward and reward-predictive cues (see **Fig. 1-2**). This may be attributable to unique dopamine receptor expression in mNAcSh, where D3-type dopamine receptors are highly enriched ^43,44^. However, we also find that mNAcSh SPNs are cue-excited but do not increase their cue-excitation across training days, and thus, do not encode reward expectation as is canonically observed in Nac core ^13,15^. DA is also likely not released as robustly to reward-predictive cues in mNAcSh as it is in Nac core ^18,45^. Recent evidence also demonstrates that evoked DA release in mNAcSh does not reinforce simultaneously presented cues ^17^. Altogether, these converging lines of research and the present study suggest that DA is not a major mechanism underlying cue information processing and cue reinforcement in mNAcSh, like it is in Nac core.

Mu opioid receptor (MOPR) activation in the basal forebrain has been thoroughly characterized for its localized robust induction of reward ‘liking’ and ‘disliking’. In particular, MOPR activation in anterior VP suppresses reward ‘liking’ and consumption while MOPR activation in posterior VP enhances reward ‘liking’ and consumption ^46,47^. Penk-SPNs in mNAcSh likely release enkephalin, which is a high affinity endogenous ligand for MOPR ^48^. Here we find that mNAcSh Penk-SPNs densely project to VP and are activated by lower preference rewards. Enkephalinergic activation of MOPR from Penk-SPNs may be a potential mechanism by which reward preference signals are transduced in VP. This potential opioidergic mechanism is beyond the scope of the present study but presents an interesting avenue for further investigation.

Although one can imagine mant future studies which are necessary to understand the precise neurotransmitter and circuit-level mechanisms underlying reward identity, preference and value computation in the NAc-VP reward system, the results presented here provides critical framework and insight into how the mNAcSh robustly represents distinct reward identities, relative preference and transmits information through pallidal efferents. This neural encoding mechanism in NAc like acts in a centralized manner to integrate incoming reward information to drive motivated and consummatory behaviors.

## Methods

### Animals

Adult C57BL/6 (18–35 g) male and female wildtype, prodynorphin-IRES-Cre (*dyn*-cre) and preproenkephalin-IRES-Cre (*enk*-cre) mice were group housed, given access to food pellets and water ad libitum, and maintained on a 12 hr:12 hr light:dark cycle. All mice were kept in a sound-attenuated, isolated holding facility one week prior to surgery, post-surgery, and throughout the duration of the behavioral assays to minimize stress. For cell-type specific optogenetic experiments, we used Cre-cage and littermate controls. Unless otherwise noted, animals had *ad libitum* access to food and water. The mice were bred at Washington University in Saint Louis or the University of Washington. All animals were initially test naive, individually assigned to specific experiments as described, and not involved with other experimental procedures. Statistical comparisons did not detect any significant differences between male and female mice and were therefore combined to complete final group sizes. All animals were monitored for health status daily and before experimentation for the entirety of the study. All procedures were approved by the Animal Care and Use Committee of Washington University, Animal Care and Use Committee of the University of Washington and conformed to US National Institutes of Health guidelines.

### Tissue collection after behavioral experiments

Unless otherwise stated, animals were transcardially perfused with 0.1 M phosphate-buffered saline (PBS) and then 40 mL 4% paraformaldehyde (PFA). Brains were dissected and post-fixed in 4% PFA overnight and then transferred to 30% sucrose solution for cryoprotection. Brains were sectioned at 30 mM on a microtome and stored in a 0.01M phosphate buffer at 4°C prior to immunohistochemistry and tracing experiments. For behavioral cohorts, viral expression and optical fiber placements were confirmed before inclusion in the presented datasets.

### RNAscope Fluorescent *In Situ* Hybridization

Following rapid decapitation of wildtype mice, brains were rapidly frozen in 100mL -50°C isopentane and stored at -80°C. Coronal sections corresponding to the site of interest or injection plane used in the behavioral experiments were cut at 20uM at -20°C and thaw-mounted onto SuperFrost Plus slides (Fisher). Slides were stored at -80°C until further processing. Fluorescent in situ hybridization was performed according to the RNAscope 2.0 Fluorescent Multiplex Kit User Manual for Fresh Frozen Tissue (Advanced Cell Diagnostics, Inc.). Briefly, sections were fixed in 4% PFA, dehydrated, and treated with pretreatment 4 protease solution. Sections were then incubated for target probes for mouse proenkephalin (*Penk*, accession number <u>NM_001002927.2</u>, probe region 106 – 1332) or prodynorphin (*Pdyn*, accession number <u>NM_018863.3</u>, probe region 33 - 700). All target probes consisted of 20 ZZ oligonucleotides and were obtained from Advanced Cell Diagnostics. Following probe hybridization, sections underwent a series of probe signal amplification steps followed by incubation of fluorescently labeled robes designed to target the specific channel associated with the probes. Slides were counterstained with DAPI, and coverslips were mounted with Vectashield Hard Set mounting medium (Vector Laboratories). Images were obtained on an Olympus Fluoview 3000 confocal microscope and analyzed with HALO software. To analyze the images, each image was opened in the HALO software. DAPI positive cells were then registered and used as markers for individual cells. An observer blind to the brain tissue origin and probes used then counted the total number of probe-labeled cells for each channel separately. A positive cell consisted of an area within the radius of a DAPI nuclear staining that measured at least 10 total positive pixels for neurotransmitter probes. Two - three separate slices from the NAc or were used for each animal and that total is presented in the data.

### Stereotaxic Surgery

After mice were acclimated to the holding facility for at least seven days, the mice were anaesthetized in an induction chamber (1%-4% isoflurane) and placed into a stereotaxic frame (Kopf Instruments, model 1900) where they were mainlined at 1%-2% isoflurane. A small hole was drilled in the skull above the target brain region. For mice receiving viral injections, a needle syringe (Hamilton Company, 65458) was used to deliver the vector at a rate of 100 μL/min. Animals used for 2-photon calcium imaging experiments were injected unilaterally with 500 uL of AAVdj-ef1a-DIO-GCaMP6s into the mNAcSh (coordinates: AP +1.3, ML +0.5, DV -4.4 from bregma), implanted with a GRIN lens (Inscopix, 7.3 mm length, 0.6 mm diameter) and had a 1 cm diameter headring horizontally adhered to the top of the skull. Animals used for fiber photometry experiments were injected unilaterally with 400 uL of AAVdj-ef1a-DIO-GCaMP6s into the mNAcSh (coordinates: AP +1.3, ML +0.5, DV -4.4 from bregma), implanted ipsilaterally with fiber optic cannulas (Doric Lenses, 400 um diameter, 6 mm length, 0.48 NA) in VP (coordinates: AP +0.3, ML +1.2, DV -4.7 from bregma), LH (coordinates: AP -1.3, ML +1.1, DV -4.7 from bregma) or VTA (coordinates: AP -3.3, ML +0.4, DV -4.4 from bregma), and had a 1 cm diameter headring horizontally adhered to the top of the skull. Animals used for retrotracing and FISH were injected unilaterally with 200 uL of green or red retrobeads (Lumafluor, Retrobeads IX) in VP (coordinates: AP +0.3, ML +1.2, DV -4.7 from bregma), LH (coordinates: AP -1.3, ML +1.1, DV - 4.7 from bregma) or VTA (coordinates: AP -3.3, ML +0.4, DV -4.4 from bregma). Animals used for optogenetics experiments were injected bilaterally with 200 uL of AAV5-DIO-ChR2-EYFP or AAV5-DIO-parapinopsin-Venus into the mNAcSh (coordinates: AP +1.3, ML +/-0.5, DV -4.4 from bregma), implanted bilaterally with custom-built fiber optic cannulas (200 um diameter) in VP (coordinates: AP +0.3, ML +1.2, DV -4.7 from bregma) or LH (coordinates: AP -1.3, ML +1.1, DV -4.7 from bregma), and had a 1 cm diameter headring horizontally adhered to the top of the skull. All implants and headrings were secured to the skull using dental cement (Parkell, C&B Metabond). Animals were injected with slow-release Carprofen (5 mg/kg) prior to surgery to act as a post-surgery analgesic. Animals were monitored daily for 7 days following surgery.

### Water Access Restriction prior to behavioral testing

Mice were allowed to recover from surgery for a minimum of 4 weeks prior to water restriction or behavioral testing. To adequately motivate consistent sucrose consumption in behavioral subjects, animals were water-restricted by giving them access to water only once per day as previously described ^5,20,21^. Starting 5 days prior to behavioral testing animals were given 1.0 ml of water in a 2x2cm paper tray in their home cage each day at approximately 5-6pm, after behavioral testing. Animals readily consumed the water while supervised by the researcher to ensure they consumed most of the presented amount. Animals were also weighed routinely to ensure they stayed at 85-90% of their original body weight while under water-restriction. No health problems related to dehydration arose throughout the water-restriction period of up to ∼16 days. After the conclusion of a behavioral testing schedule, water-restriction was immediately stopped and animals were given ad-libitum access to water in their home cages again.

### Brief Access Two Bottle Choice

To test the relative preference of the 4 variable sucrose reward conditions (0%, 3%, 10%, 30% sucrose dissolved in water) while minimizing effects from satiety, we tested mice in an adapted 2-bottle choice assay where the animals were only given brief access (20 s) to sucrose sippers with a variable intertrial interval (20-40s) between access periods. Brief access 2-bottle choice assays were conducted within enclosed operant chambers, that had two retractable sucrose sippers side-by-side (MedAssociates, MED-307A, ENV-352). Discrete licks on the metallic sippers were recorded by a contact licometer system (MedAssociates, ENV-250). Prior to 2-bottle choice test days, each animal was trained with free access (2 days) and variable access (2 days) to both sippers containing 10% sucrose for 30 minute sessions. Each preference test involved presentation of 2 of the 4 different sucrose conditions and each session had two phases. In the first phase, animals were given access to either sipper in isolation so that they could sample them independently. In the second phase, animals were simultaneously presented with both sippers side-by-side and had to choose where to allocate their sipper access time. The 6 possible pairings of the 4 sucrose reward conditions were presented in a pseudorandomized and counterbalanced sequence to minimize effects from condition history and right versus left sipper location bias. Preference test days were also buffered by 10% versus 10% sucrose days to neutralize any left-versus-right biases that arose from test days.

As indicated in the Results, some sessions included a variation of this paradigm where one of the sucrose sippers contained no liquid and instead triggered closed loop optogenetic stimulation. Licking the empty sucrose spout would register a lick in the MED-PC program and immediately trigger a TTL sent from the MED-307A to an Arduino Uno. The Arduino Uno controlled a 470 nm light laser (4 mW intensity at end of patch cable, 10 Hz, 10 ms pulse width, 2 second in duration) that was optically coupled to the fiber implants on the animal’s head.

### Variable Sucrose Access during Reversible Head-fixation

After 4-6 weeks of recovery from surgery, animals were water restricted (see above) and trained to sip 10% sucrose reward at variable intervals while reversibly immobilized and head-restrained. Reversible head-fixation was conducted by scruffing the animals, backing them into a 50 ml canonical tube and sliding the animals and tube into place in a custom headstage as has been previously described ^20,21^. All animals displayed no signs of physical distress and consistent licking behavior while immobilized after 4 days of lick training and were then tested in 30 minute, 20 trial sessions each day for 4 consecutive days. Intertrial interval was uniformly distributed between 70 and 100 seconds. On each trial, animals were presented with 5 droplets of sucrose reward with a total aggregate volume of 10 ul, with 1 second between droplet delivery. Reward delivery was controlled by custom code on an Arduino Uno. Animals immediately licked the presented sucrose reward, with an average licking bout duration of 7-8 seconds. In each session, animals consumed 1 of 4 possible sucrose reward conditions (0%, 3%, 10% or 30% sucrose in water), with the order of condition days pseudorandomized and counterbalanced across animals. Discrete lick events were detected and recorded with <1 ms time resolution using a contact licometer circuit as has been previously described ^20,21^. In some cases, 15-17 FPS behavioral video of the animal’s face was recorded alongside the contact licometer or instead of the contact licometer. The presence of sucrose licking in a particular frame was assessed manually or through automated scoring by a recurrent neural network model (see below) that was extensively trained and validated using simultaneous contact licometer information. The behavioral testing was conducted in dark or far-red light conditions to remove contamination from visual stimuli. 2-photon calcium imaging data or photometry data was acquired concurrently with this behavioral paradigm on days 1-4.

### Pavlovian conditioning during Reversible Head-fixation

The same animals that were tested in the variable sucrose during reversible head-fixation paradigm (see above) were then also trained to associate an auditory cue to subsequent delivery of 10% sucrose reward across 7 days of Pavlovian conditioning during reversible head-fixation. Animals were again tested in 30 minute, 20 trial sessions each day for 7 consecutive days. Intertrial interval was uniformly distributed between 70 and 100 seconds. On each trial, animals were presented with 3 seconds of a 4 kHz auditory cue. 2 seconds after cue termination, 10% sucrose reward was delivered in a series of 5 droplets with total aggregate volume of 10 ul, with 1 second between droplet delivery. Reward delivery and cues were controlled by custom code on an Arduino Uno. The latter half of trials (last 10 out of 20 trials) on Pavlovian day 7 were reward omission trials, where the cue was presented as usual, but the expected sucrose reward was not delivered. Licking behavior was detected and recorded with <1 ms time resolution using a contact licometer and/or by 15-17 FPS behavioral video. The behavioral testing was conducted in dark or far-red light conditions to remove contamination from visual stimuli. 2-photon calcium imaging data or photometry data was acquired concurrently with this behavioral paradigm on Pavlovian training days 1, 3, 5 and 7.

### Automated Lick Detection from Video using Recurrent Neural Network

In some reversible head-fixed behavior sessions, 15-17 FPS, 720p or 1080p behavioral video was collected concurrently with or instead of the contact licometer data for lick detection. The camera was positioned approximately 6-8 cm in front of the animal’s face to capture licking behavior. Far-red or infrared light was used to illuminate the animals face during behavioral video acquisition. ROIs near the animal’s mouth (160x160 pixels for 1080p video, 100x100 pixels for 720p video) were manually selected and cropped from the original video. Pixel data types were converted from 16-bit RGB to 8-bit integer grayscale. Individual pixels were normalized by taking the absolute value of the z score, using the mean and standard deviation of an individual pixel over the duration of the entire video. Normalized frames were then spatially downsampled to 5x5 pixels. 48 behavioral sessions that had simultaneous contact licometer lick detection and behavioral video were used to identify which video frames contained spout licks. 10 frame sequences of these ground-truth labelled video frames were then used to train a custom recurrent neural network (RNN) (10 long-short-term-memory node layer, followed by 10 node dense layer) using standard tools from the Python TensorFlow library. A RNN implementation was chosen because it considers changes in pixels across sequential frames, rather than only considering one frame at a time. The RNN model for binary frame classification performed with high accuracy and specificity with a ROC AUC = 0.91. The model was then used to predict licking for every frame of 104 30-minute behavioral videos. To further validate the accuracy of the automated detection approach, the predicted classifications of the model were compared to manually scored video data where researchers scored 1 second time bins as containing animal licks or not. The model again reliably matched the compared lick scoring information, with 82% median overlap between manually scored bins and recurrent neural network frames (see **Extended Data Fig. 5**). Furthermore, the overall behavioral findings of how much animals licked for different reward conditions and to reward-predictive cues across Pavlovian conditioning were consistent between contact licometer, RNN-scored video and manually-scored video approaches for lick detection. Code for RNN lick detection model is available at github.com/christianepedersen.

### In vivo 2-photon Calcium Imaging Acquisition

Image acquisition was performed simultaneously with reversible head-fixed behavior (see above) using the Olympus Fluoview FVMPE-RS 2-photon microscope. A 20x magnification air objective lens (Olympus, LCPLN20XIR, 0.45 numerical aperture, 8.3mm working distance) was optically coupled to the surgically implanted endoscopic lens (Inscopix, 7.3mm length, 0.6mm diameter GRIN lens) of an animal. 920 nm laser light (SpectraPhysics MaiTai Laser, ∼100-femtosecond pulse width) excited GCaMP6s fluorophores during resonant scanning (30-Hz roundtrip frame rate, averaging every 6 frames in real-time). Emitted ∼535 nm photons were directed to detectors (Olympus, GaAsP PMT) and attributed to the excited pixel. The same focal plane (<5 um tolerance) was optically accessed for each imaging session with the same animal so we could reliably track the activity of the same neurons across multiple days and conditions. A 200-frame, 40-second image series was initiated 20 seconds before reward delivery on each behavioral trial and was triggered by an Arduino Uno that also controlled reward delivery and cues.

### *In vivo* 2-photon Calcium Imaging Data Processing

The Olympus OIR files collected during imaging through Olympus FluoView (FV1200) were exported as tif files. Tif files for individual behavioral trials were motion corrected using a nonrigid motion correction package (Flatiron Institute: github.com/flatironinstitute/NoRMCorre) and concatenated using custom code in MatLab. These whole-session tif files were then converted to an HDF5 format using a custom code. These HDF5 files were then motion corrected in the x–y plane using an adapted hidden Markov model (SIMA version 1.3.0: losonczylab.org/sima/1.3/api/motion.html) as previously reported ^49^. We had found that the imaging plane showed very little z movement in mNAcSh (<5 µm based on a random sample of sessions). Individual neurons were then identified manually and selected by drawing regions of interest (ROIs) manually using the ImageJ polygon tool on the mean projection of pixel intensity across all frames in a session. ROIs for different sessions that corresponded to the same neuron and accurately captured the dynamics of the cell were used to ‘track’ that neuron across days and conditions. Individual neurons that did not have consistent ROIs for all sessions were not ‘tracked’ and were not used for comparison of responses to different conditions. The retained ROIs were imported into SIMA and then used for signal extraction.

Custom code added neuropil correction to the SIMA signal extraction as previously described ^21^. GCaMP6s labelled neurons were fairly sparse in the focal plane, with minimal overlap of neurons in x, y dimensions. However, changes in the fluorescence of individual neuron ROIs were still contaminated with slight variation in back fluorescence due to neuropil. Neuropil correction was done by first calculating a neuropil signal around each ROI. This was done by calculating a weighted sum of all recorded pixels excluding those falling within a 15-pixel (∼17-µm) radius of all ROIs. The weight for any pixel was calculated using a Gaussian function centered on the ROI of interest with a radius of 50 pixels (∼45µm). These parameters were obtained after a systematic search of the parameter space in a small subset of sessions and visually comparing the obtained fluorescence traces against the raw videos in the current mNAcSh dataset as well as in previous studies ^21^. The results in this manuscript are robust against large variation in these parameters. Once the neuropil signal was calculated for every ROI, a correction of this signal was done by subtracting 0.8 multiplied by this signal from the raw calcium trace of the ROI. The results of this previously published neuropil correction procedure ^21^ were also found to be highly consistent with results from another prominent background subtraction approach from the package Suite2p ^50^. After single cell fluorescence time series were extracted and neuropil-corrected from raw calcium image sequences, they were detrended by fitting a double exponential curve to the entire session and subtracting it. Changes in fluorescence from the now consistent baseline were z-score normalized, using the mean and standard deviation of the entire session.

### *In Vivo* 2-photon Calcium Imaging Data Analysis

Individual neurons that were successfully tracked across days and conditions were further analyzed to understand their responsivity to different behavioral conditions. First, the average response of individual neurons to all variable sucrose and Pavlovian conditions was taken by taking the mean fluorescence of each cell across all 20 trials of a given behavioral session (or 10 rewarded trials or 10 extinction trials for Pavlovian Day 7). Prior to clustering, dimensionality reduction was performed on trial-averaged responses to all reward conditions (0-10 seconds from reward onset for variable sucrose sessions, -5-10 seconds from reward onset for Pavlovian sessions) to capture the most important Euclidean distance measures during subsequent clustering. Specifically, principal component analysis was performed and the number of components retained (6-8 components) was chosen through standard Scree methods. The principal components of reward responses of individual neurons across all variable sucrose or Pavlovian conditions were then used for K-means clustering (number of clusters with lowest silhouette score was chosen, 500x bootstrapping was performed to minimize variability from initial conditions) to group together neurons that had similar responses across these different conditions. K-means clustering and silhouette score calculations were performed in Python using the Scikit-learn library. Several clusters were identified among dyn-SPN and enk-SPN neurons for each behavioral paradigm. The trial-averaged responses for all neurons within a given cluster were then further averaged to generate the average response traces for each cluster to every condition.

To investigate whether the different reward conditions in the variable sucrose paradigm (0%, 3%, 10%, 30%) could be uniquely identified only by the simultaneous activity of mNAcSh SPNs, a multi-class linear support vector classifier (SVC) model (cost = 0.8) was created. The linear SVC model was trained and tested on the data for each animal individually. The class of the reward condition and the magnitude of the reward response (0-8 s from onset of reward delivery) of every neuron for a given trial was used as input for the model (80 trials per animal, 20 trials per reward class). 80% of the trials were used to train the linear SVC and 20% were used for cross-validation. The accuracy of the model during cross-validation for each animal is reported in the Results. The linear SVC model was created in Python using the Scikit-learn library and feature coefficients were taken directly from the model and averaged by which K-means cluster the neurons belonged to. To test whether cue-excitation of mNAcSh SPNs was correlated to Pavlovian conditioning session index, trial index within a session or to anticipatory licking to the cue, Pearson’s correlations were calculated relative to the magnitude of cue-response (0-5 s from onset of cue) of SPNs that belonged to clusters that were both cue-excited and reward-excited.

To investigate the relationship between SPN cue and reward-response and relative anatomical position, we calculated the relative position of tracked SPNs in the focal plane during 2-photon calcium imaging sessions. The 2-photon resonant scanner generated raw frames that were 512x512 pixels. The x and y pixel position at the center of each ROI that was manually drawn in ImageJ was considered for each successfully tracked neuron. The relative anatomical position of each neuron was calculated as the mean x and y position of its ROI centers across all imaging sessions. Medial-lateral and anterior-posterior axes were inferred based on the orientation of the animal being orthogonal to the objective lens during the imaging and behavior session. The depth of the focal plane ventral to the GRIN lens during the imaging session introduces a complex magnification effect, so the precise ratio between pixels and anatomical distance in the brain could only be approximated based on the 600 um diameter of the lens and the width of the focal plane being approximately equivalent.

### *In Vivo* Fiber Photometry During Reversible Head-fixation

Fiber photometry recordings were made throughout the entirety of 20 trial, 30-minute head-fixed variable sucrose and Pavlovian conditioning sessions using a previously published approach ^51^. Prior to recording, an optic fiber was attached to the implanted fiber using a ferrule sleeve (Doric, ZR_2.5). Two LEDs were used to excite GCaMP6s. A 531-Hz sinusoidal LED light (Thorlabs, LED light: M470F3; LED driver: DC4104) was bandpass filtered (470 ± 20 nm, Doric, FMC4) to excite GCaMP6s and evoke Ca^2+^-dependent emission. A 211-Hz sinusoidal LED light (Thorlabs, LED light: M405FP1; LED driver: DC4104) was bandpass filtered (405 ± 10 nm, Doric, FMC4) to excite GCaMP6s and evoke Ca^2+^-independent isosbestic control emission. Prior to behavior and recording, a 120 s period of GCaMP6s excitation with 405 nm and 470 nm light was used to remove the majority of baseline drift. Laser intensity for the 470 nm and 405 nm wavelength bands were measured at the tip of the optic fiber and adjusted to ∼70 μW before each day of recording. GCaMP6s fluorescence traveled through the same optic fiber before being bandpass filtered (525 ± 25 nm, Doric, FMC4), transduced by a femtowatt silicon photoreceiver (Newport, 2151) and recorded by a real-time processor (TDT, RZ5P). The envelopes of the 531-Hz and 211-Hz signals were extracted in real-time by the TDT program Synapse at a sampling rate of 1017.25 Hz.

### Photometry Data Analysis

Custom MATLAB scripts were developed for analyzing fiber photometry data in context of mouse behavior and can be accessed online (github.com/christianepedersen). The isosbestic 405 nm excitation control signal was subtracted from the 470 nm excitation signal to remove movement artifacts from intracellular Ca^2+^-dependent GCaMP6s fluorescence (see **Extended Data Fig. 10**). Baseline drift was evident in the signal due to slow photobleaching artifacts, particularly during the first several minutes of each 30-minute recording session. A double exponential curve was fit to the raw trace and subtracted to correct for baseline drift. After baseline correction, the photometry trace was z-scored relative to the mean and standard deviation of the entire session. The processed fluorescence timeseries was then aligned to animal behavior and the average fluorescence relative to the time of behavioral events (sucrose reward delivery) was visualized. Quantification of peri-event fluorescence was calculated by taking the average of all samples over a given time window for each trial and comparing between behavioral conditions.

### Optogenetic Stimulation during Head-fixed Sucrose Consumption

After recovering from surgery, animals were water-restricted and trained to lick 10% sucrose reward while reversibly immobilized and head-restrained. After 4 days of lick training, all animals displayed consistent lick behavior in response to variable interval sucrose presentation and were then put through 2 consecutive test days. Each test day involved one 30 minute, 20 trial session in which the animals were presented with either 10% or 30% sucrose reward. The order of sucrose condition days was pseudorandomized and counterbalanced across animals. For consistency with the imaging and photometry behavior paradigms, animals were presented with 5 droplets of sucrose reward per trial with a total aggregate volume of 10 ul and 1 second between droplet delivery. Animals immediately licked the presented sucrose reward, with average licking bout durations of 7-8 seconds. On a pseudorandom 50% of the trials within a session, time-locked optogenetic manipulation of mNAcSh-VP^enk^ or mNAcSh-LH^dyn^ activity would be paired with reward delivery, via 4 seconds of 470 nm blue laser light delivery (4 mW for ChR2; 10 mW for PPO measured at end of patch cable, 10 Hz, 10 ms pulse width). 10Hz optogenetic stimulation frequency was chosen because it is physiologically relevant level of neural activity for SPNs during phasic reward-related activity ^48^. Any differences in consummatory licking between no laser and laser trials were recorded via the contact licometer circuit with <1 ms time precision. All differences in licking between no laser and laser trials were evaluated within subject to account for the baseline variability in lick rate between animals. To control for possible behavioral effects from the delivery of the blue light itself, cre-littermate controls were also injected, implanted, and tested in identical conditions.

### Open-field Locomotion Assay

To determine whether mNAcSh-VP^Penk^ or mNAcSh-LH^Pdyn^ is naturally activated during bouts of motor activity as they are during reward consumption (see **Fig. 6, Extended Data Fig. 9**), photometry animals were tested in a locomotion assay. Animals freely roamed a 0.5x0.5 m arena in far red light conditions in a quiet environment to measure the amount of spontaneous locomotion in a context that was not anxiogenic. Simultaneous fiber photometry recordings were conducted during the 30-minute long behavioral sessions. 15 FPS video was captured throughout the behavioral session by a camera position directly above the behavioral arena. Automated animal location tracking software (Noldus, Ethovision) was used on the video data after the session to get the x and y coordinates of the animal in the behavioral arena for every frame of the video. Changes in animal coordinates between frames were used to calculate the instantaneous velocity of the animal for each frame. Pearson’s correlations were calculated between the velocity of the animal and z scored photometry fluorescence for every frame. Local maxima for movement velocity were identified as discrete events by using the MatLab ‘findpeaks()’ command (peak prominence >1 SD, peak width >4 seconds). Windows of the fluorescence time series were aligned to velocity peaks to visualize peri-velocity peak averaged fluorescence. Contrary to the mNAcSh-VP^Penk^ and mNAcSh-LH^Pdyn^ reward-excitations observed during sucrose consumption, bouts of substantial locomotion did not correlate to circuit recruitment.

To determine if mNAcSh-VP^Penk^ or mNAcSh-LH^Pdyn^ activity manipulation could causally disrupt the execution of motor actions, we then performed a similar locomotion assay experiment with intermittent optogenetic photo-excitation or photo-inhibition. Bilaterally implanted optogenetics animals (see above) freely roamed a 0.5x0.5 m arena in far red light conditions in a quiet environment to measure the amount of spontaneous locomotion in a context that was not anxiogenic. Every minute of the 14 minute behavioral session had 30 seconds of pulsing 470 nm laser light (4 mW for ChR2; 10 mW for PPO measured at end of patch cable, 10 Hz, 10 ms pulse width) followed by 30 seconds of no laser. 15 FPS behavioral video was captured throughout the session and was again used to calculate the instantaneous velocity of the animal for every frame. Movement velocity was compared within subject and within session between laser on epochs and laser off epochs. Optogenetic photo-excitation or photo-inhibition of mNAcSh-VP^enk^ or mNAcSh-LH^dyn^ had no significant effect on animal movement velocity. There was also no cumulative effect on locomotion throughout the session as movement velocity was found to be consistent throughout the entire duration of the session.

### Statistical Approaches and Analysis

All data collected were averaged and expressed as mean ± SEM. Statistical significance was taken as ∗p < 0.05, ∗∗p < 0.01, and ∗∗∗p < 0.001, as determined by Pearson’s correlation, Student’s t-test or one-way ANOVA followed by Tukey post hoc tests as appropriate. For *in situ* hybridization data, we used Student’s t-test. For 2-photon imaging and photometry experiments, we used Pearson’s correlation and Student’s t-tests, as appropriate. For optogenetic behavioral experiments, we used pairwise t-tests for comparison within subject and within session. All n values for each experimental group are described in the appropriate figure legend. For behavioral experiments, group size ranged from n = 6 to n = 10. For *in situ* hybridization quantification experiments, slices were collected from 2-3 mice, with data averaged from 2-3 slices per mouse. Statistical analyses were performed in GraphPad Prism 8.0 (Graphpad, La Jolla, CA), MatLab 2020 (The MathWorks, Natick, MA) or with Python SciPy library.

## Acknowledgements

We thank Dylan Blumenthal, Michelle Chung, Taylor Hobbs, and Carina Pizzano for animal colony maintenance. We thank the Bruchas lab, Stuber lab and NAPE Center for helpful discussions.

## Author Contributions

C.E.P, D.C.C and M.R.B conceived the study and designed experiments. C.E.P, D.C.C., S.A.K and P.J.M. performed experiments and collected data. C.E.P analyzed data and designed analyses. M.M.G, Z.C.Z. and P.R.O contributed to experimental design and provided technical support for data collection. C.E.P., D.C.C and M.R.B wrote the manuscript.

## Data and materials availability

This study did not generate any new/unique plasmids, mouse lines or reagents. Custom MatLab and Python analysis codes were created to appropriately organize, process, and combine photometry and 2-photon recording data with associated behavioral data. Analysis code for photometry and 2-photon imaging data will be made available on github.com/christianepedersen. Data sets supporting the current study are available from the corresponding author upon reasonable request.

## Materials and Correspondence

Further information and requests for resources and reagents should be directed to and will be fulfilled by corresponding author Michael R. Bruchas (^12,13^).

---

**Extended Data Figure 1.**
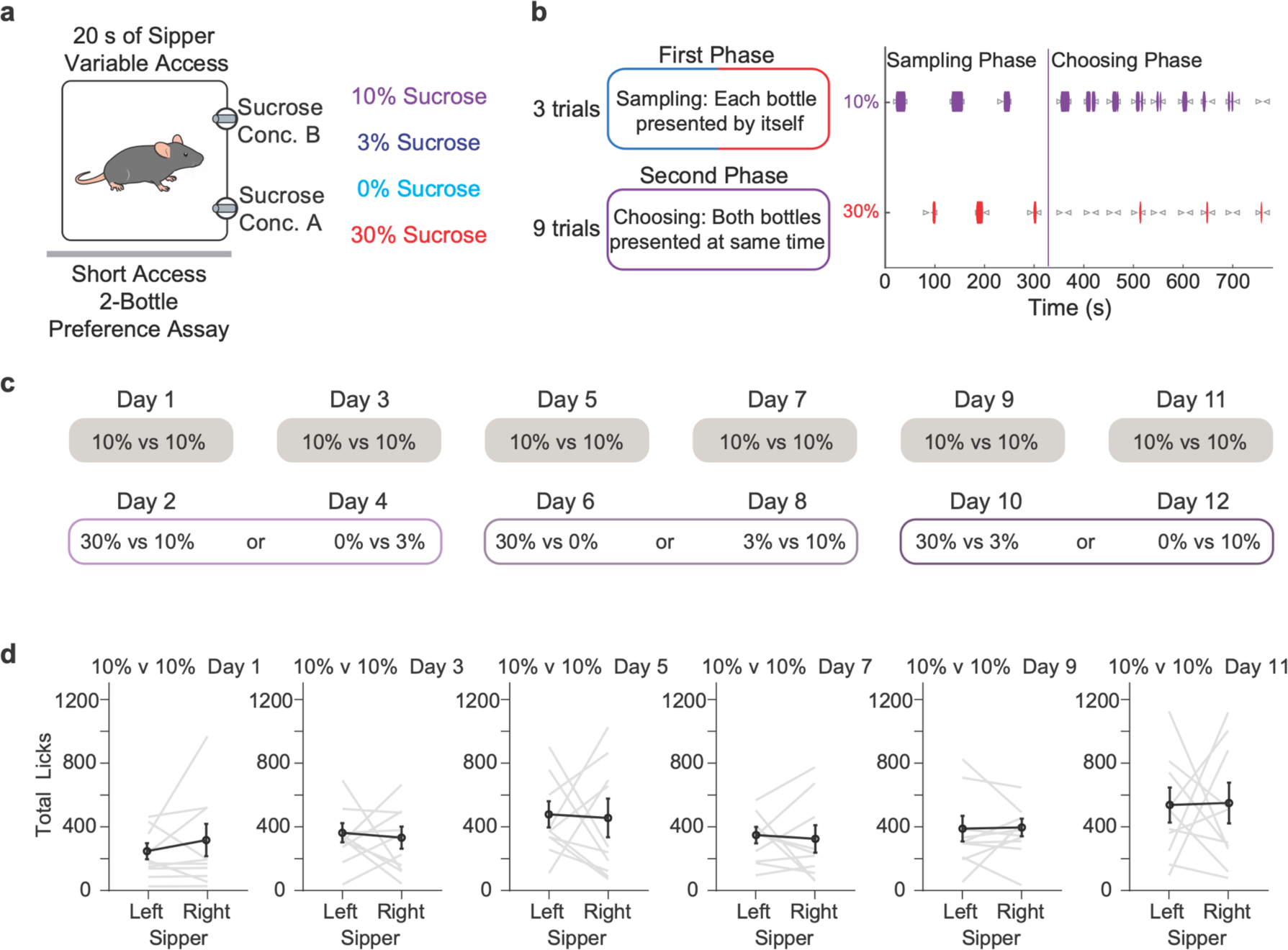
Sucrose concentration preference of C57 mice under water restriction. (a) Schematic of brief access 2-bottle choice task: animal makes choice between two sippers containing different concentrations of sucrose reward. (b) Time-course of task and representative lick data throughout session. (c) Brief access 2-bottle choice task schedule: Alternating between pseudorandomized test days and 10% versus 10% days to neutralize spatial biases that arise during test days. (d) Preference between left and right sippers containing identical rewards on spatial bias neutralization days (n = 10 animals, pairwise t-test: p = 0.411, p = 0.760, p = 0.885, p = 0.784, p = 0.911, p = 0.949).

**Extended Data Figure 2.**
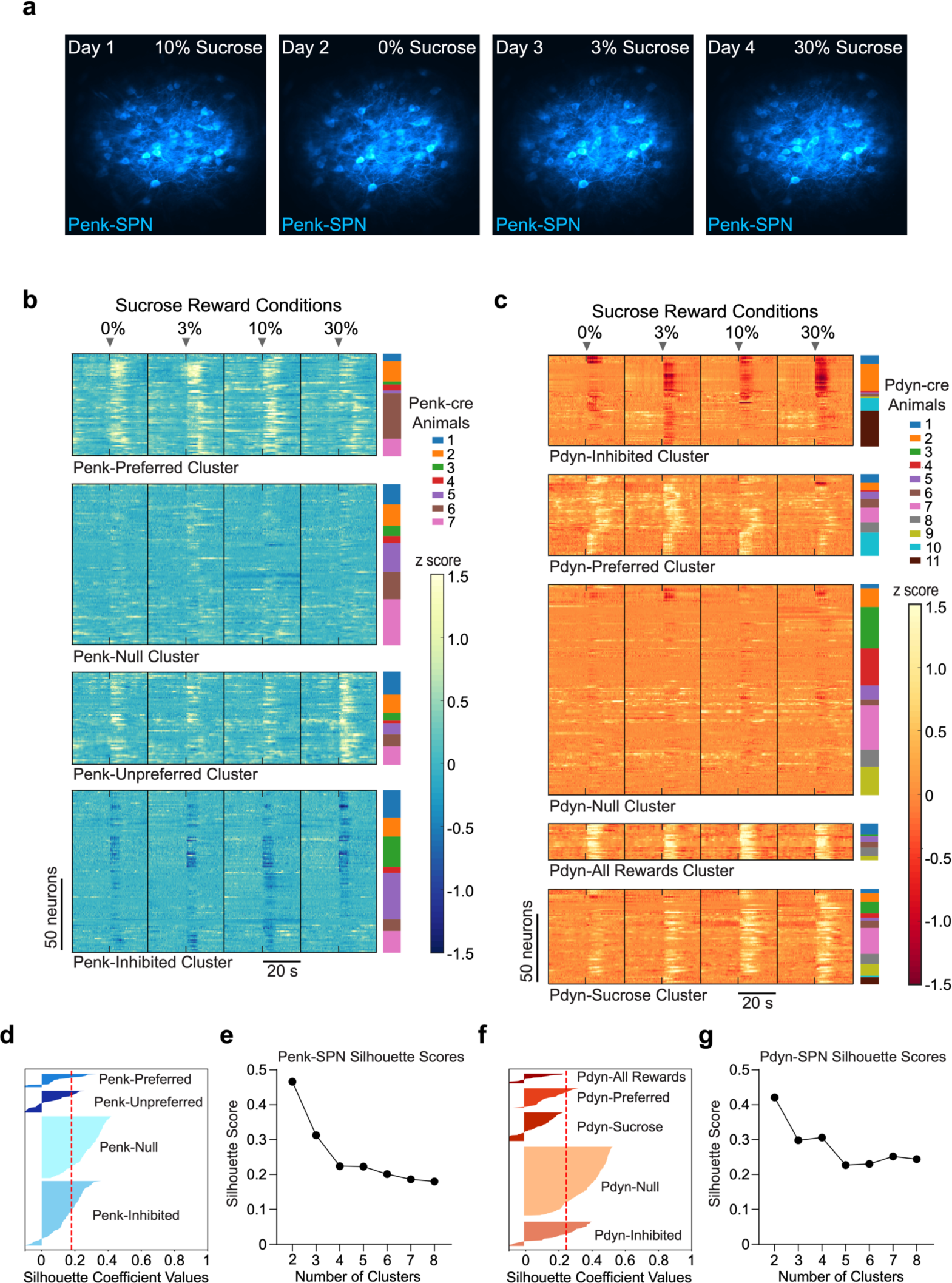
Individual SPN tracking and clustering based on response to variable sucrose conditions. (a) Mean pixel values across entire in vivo 2-photon calcium imaging session for a representative Penk-cre animal to demonstrate the ability to resolve individual cell morphology and track neurons reliably across multiple days and conditions. (b) Mean response (20 trials per condition) of individual Penk-SPNs to all sucrose concentration conditions. Individual tracked cells are arranged by their cluster ID and animal ID to show representation of different animals within the neuronal clusters. (c) Mean response (20 trials per condition) of individual Pdyn-SPNs to all sucrose concentration conditions. Individual tracked cells are arranged by their cluster ID and animal ID to show representation of different animals within the neuronal clusters. (d) Silhouette plot shown for optimal k-means clustering of Penk-SPNs. (e) Silhouette scores for different Penk-SPN cluster counts to determine the optimal number of clusters. (f) Silhouette plot shown for optimal k-means clustering of Pdyn-SPNs. (g) Silhouette scores for different Pdyn-SPN cluster counts to determine the optimal number of clusters.

**Extended Data Figure 3.**
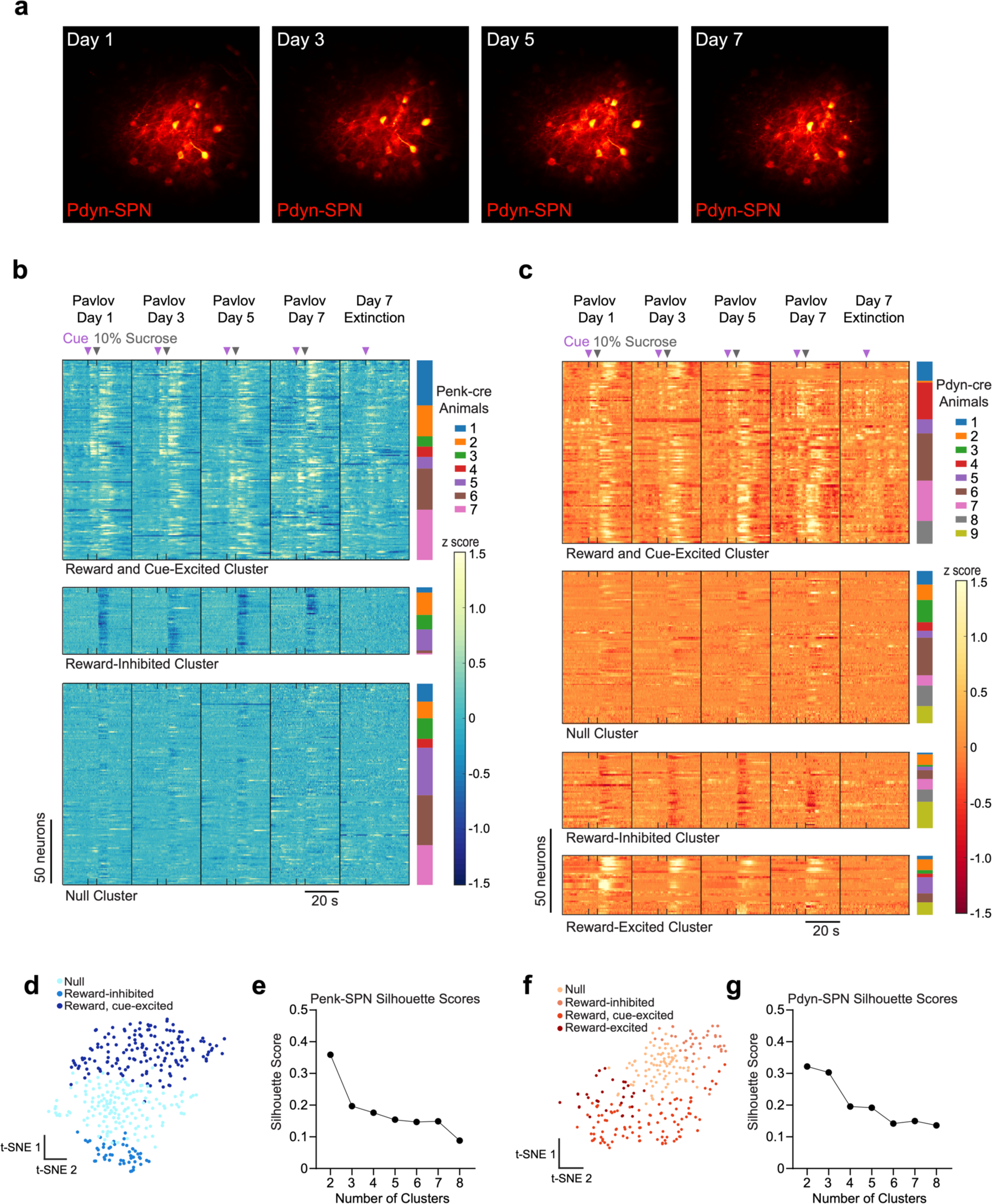
Individual SPN tracking and clustering based on response to Pavlovian conditioning. (a) Mean pixel values across entire in vivo 2-photon calcium imaging session for a representative Pdyn-cre animal to demonstrate the ability to resolve individual cell morphology and track neurons reliably across multiple days and conditions. (b) Mean response (20 trials per condition) of individual Penk-SPNs to cue and reward for all Pavlovian conditioning days. Individual tracked cells are arranged by their cluster ID and animal ID to show representation of different animals within the neuronal clusters. (c) Mean response (20 trials per condition) of individual Pdyn-SPNs to cue and reward for all Pavlovian conditioning days. Individual tracked cells are arranged by their cluster ID and animal ID to show representation of different animals within the neuronal clusters. (d) t-sne plot of variance in activity of Penk-SPN clusters during cue and reward for all Pavlovian conditioning days. Each dot represents an individual tracked neuron that composes the cluster. (e) Silhouette scores for different Penk-SPN Pavlovian response cluster counts to determine the optimal number of clusters. (f) t-sne plot of variance in activity of Pdyn-SPN clusters during cue and reward for all Pavlovian conditioning days. Each dot represents an individual tracked neuron that composes the cluster. (g) Silhouette scores for different Pdyn-SPN Pavlovian response cluster counts to determine the optimal number of clusters.

**Extended Data Figure 4.**
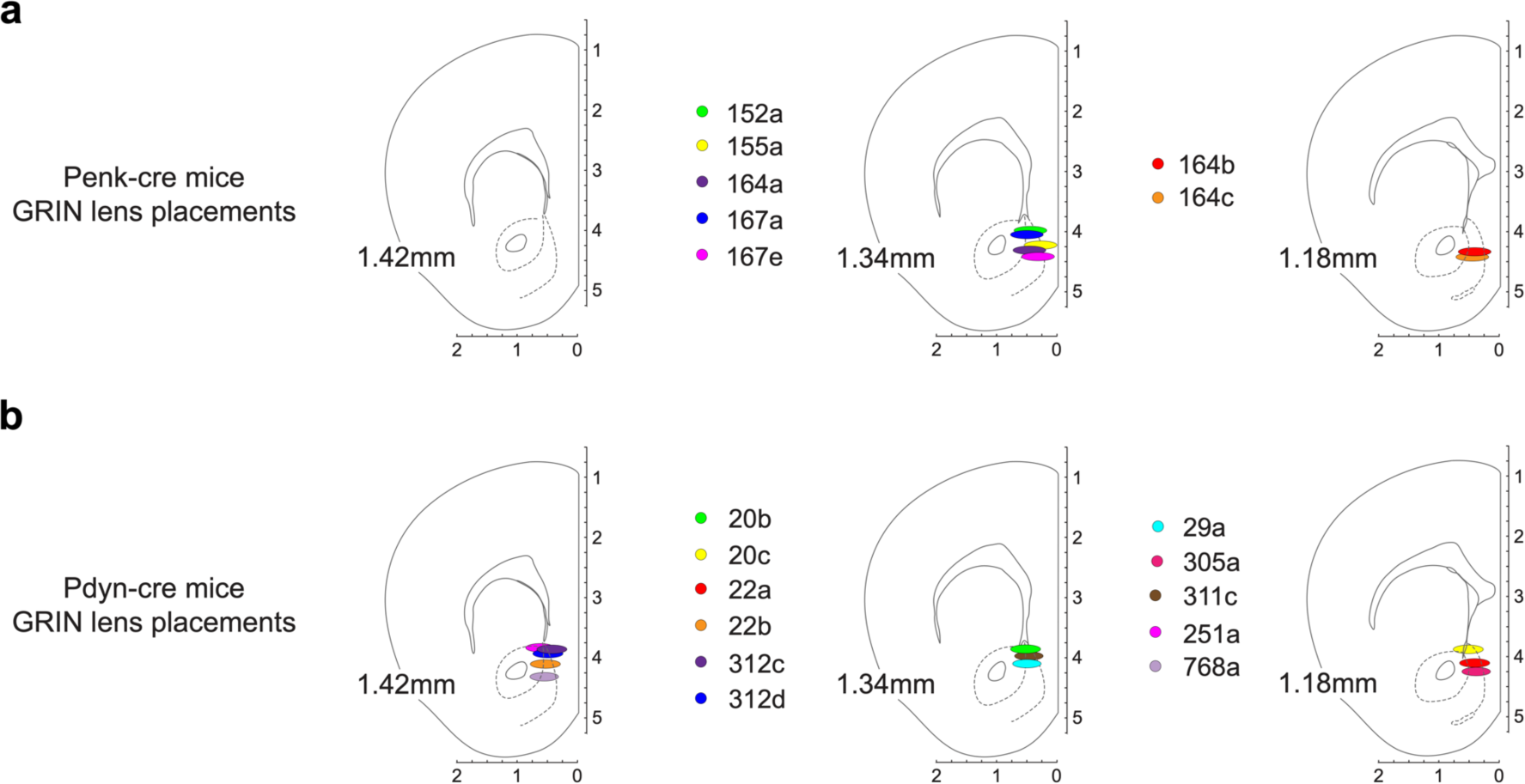
GRIN lens placements. (a) Surgical placements of endoscopic GRIN lenses in mNAcSh of Penk-cre animals. (b) Surgical placements of endoscopic GRIN lenses in mNAcSh of Pdyn-cre animals.

**Extended Data Figure 5.**
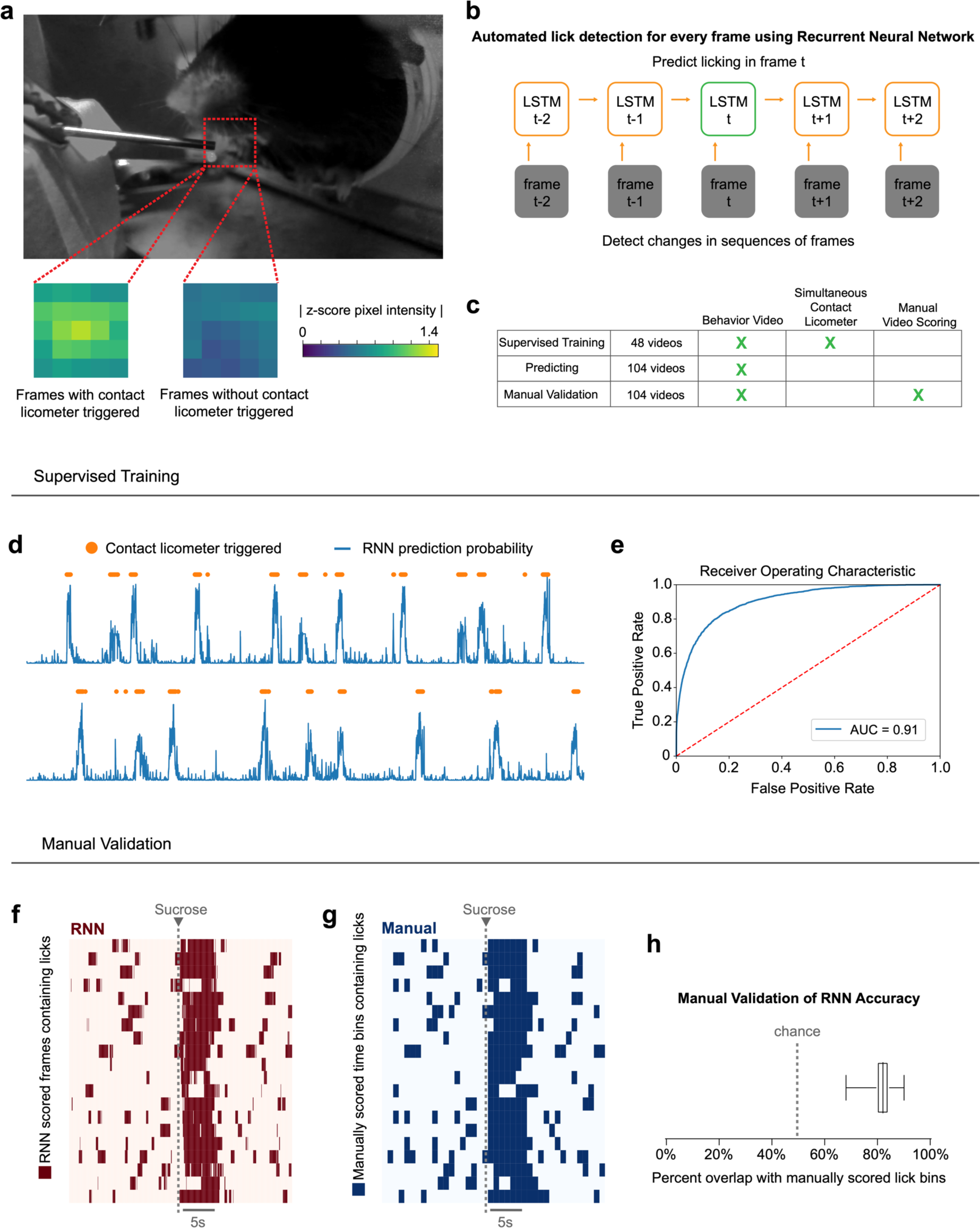
Training and Validation of Recurrent Neural Network for Lick Detection. (a) Representative frame from head-fixed behavior video. ROI near animals mouth cropped from video, spatially downsampled and normalized prior to frame classification. (b) Cartoon depicting organization of RNN that uses changes in previous and future frames to predict licking in the current frame. (c) Table depicting which data sets were used to initially train the RNN model using contact licometer information as frame labels and used to manually validate RNN model accuracy. (d) Representative sequential RNN prediction probabilities for when animal is licking (blue) overlaid with timestamps of animal licking from contact licometer (orange). (e) ROC curve showing specificity of RNN binary classification on entire 48 video training set. (f and g) Representative session frame classification by RNN model (f) and manual video scoring (g). (h) Overlap between manually scored video and RNN model video classification for all 104 behavioral videos.

**Extended Data Figure 6.**
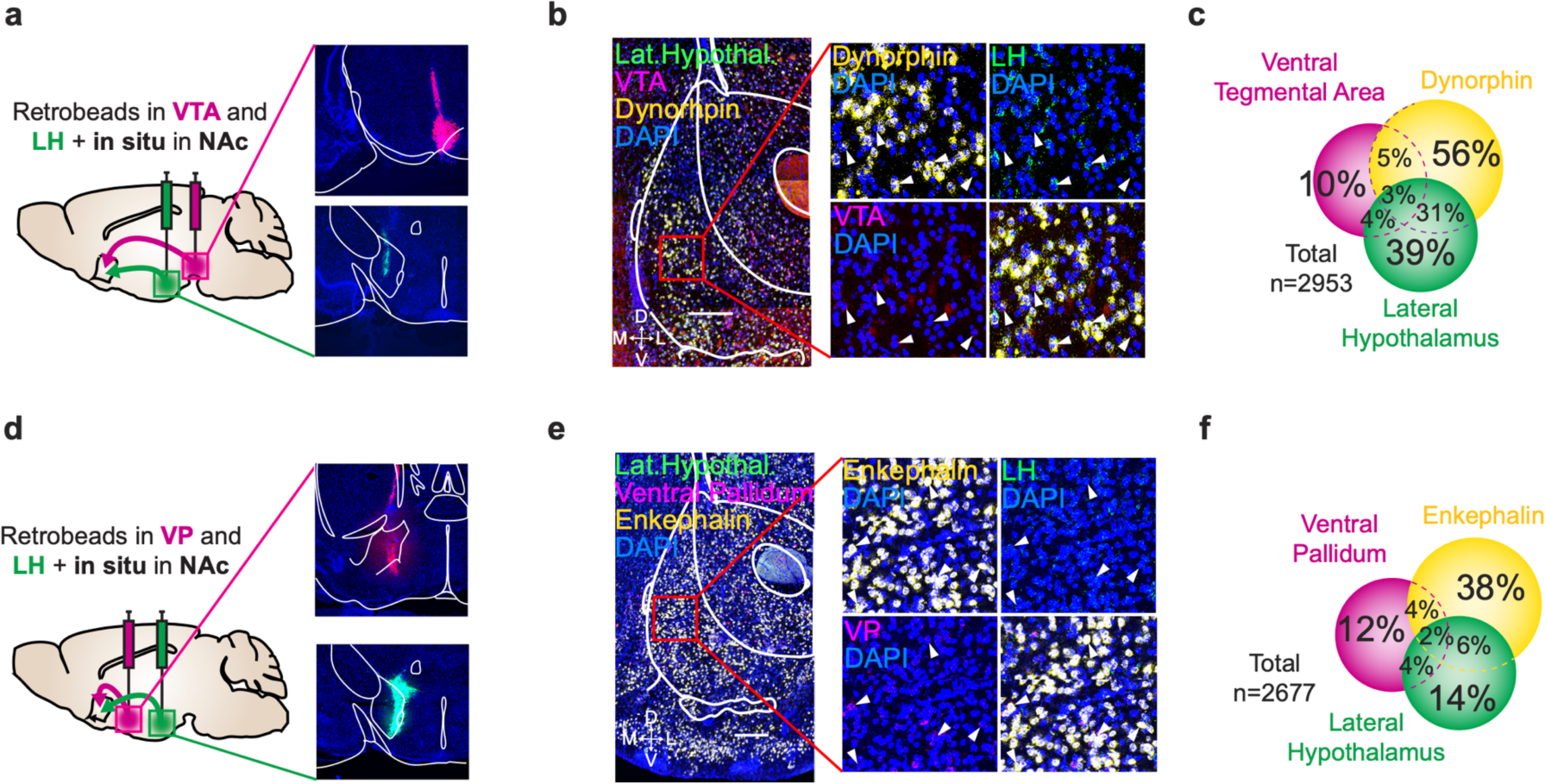
There is limited collateralization of Penk-SPN and Pdyn-SPN efferent circuits. (a) Schematic (left) and representative images (right) of ventral tegmental area (top) or lateral hypothalamus (bottom) of retrobead injection sites. (b and c) In situ hybridization (b) and quantification (c) of prodynorphin, red, and green retrobeads in mNAcSh (scale bar = 200um). Percentage of DAPI-labelled cells that are also labelled with indicated probe. (d) Schematic (left) and representative images (right) of ventral pallidum (top) or lateral hypothalamus (bottom) of retrobead injection sites. (e and f) In situ hybridization (e) and quantification (f) of proenkephalin, red, and green retrobeads in mNAcSh (scale bar = 200um). Percentage of DAPI-labelled cells that are also labelled with indicated probe.

**Extended Data Figure 7.**
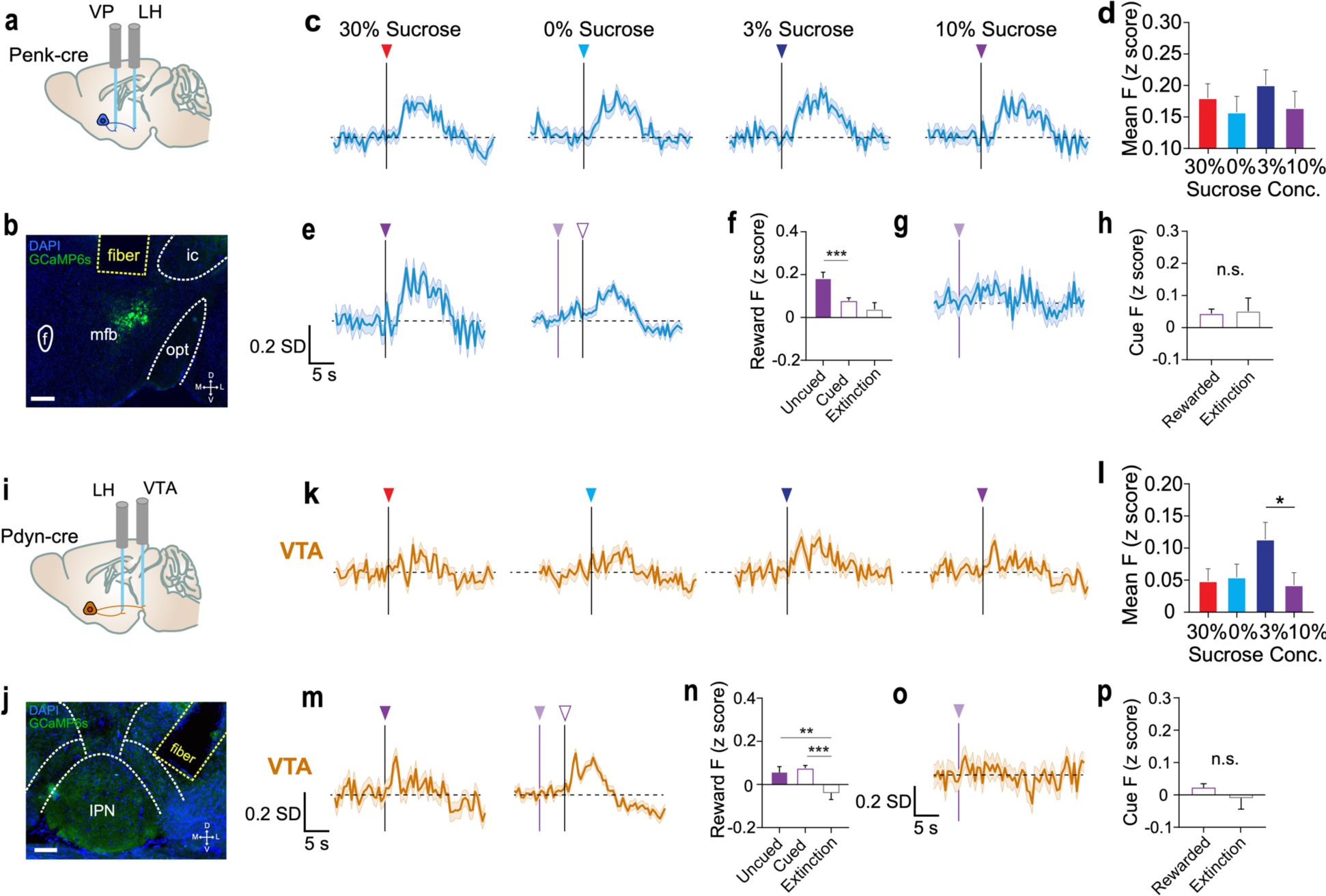
mNAcSh-LH^Pdyn^ and mNAcSh-VP^Penk^ are mildly cue-responsive. (a) Sagittal section cartoon displaying target injection region, cell-types and fiber implant locations for mNAcSh-LH^Penk^ recording. (b) Representative coronal section showing Penk-SPN terminals in LH expressing GCaMP6s underneath the fiber implant (scale bar = 200 um). (c) Mean mNAcSh-LH^Penk^ reward-response for every trial of each variable sucrose reward condition. (d) Mean mNAcSh-LH^Penk^ activity over reward response epoch (2-12 seconds from reward-onset) for each condition (n = 6 animals, 127+ trials; one-way ANOVA: p = 0.6167). (e) Mean mNAcSh-VP^Penk^ reward-response for every uncued 10% sucrose trial and every cued 10% sucrose trial. (f) Mean mNAcSh-VP^Penk^ activity over reward response epoch (2-12 seconds from reward-onset) for uncued 10% sucrose, cued 10% sucrose and extinction trials (n = 9 animals, 96+ trials; one-way ANOVA: Uncued vs Cued, *p = 0.014, Cued vs Ext., ***p < 0.001). (g) Mean mNAcSh-VP^Penk^ reward-response for every trial of day 7 Pavlovian extinction. (h) Mean mNAcSh-VP^Penk^ activity over cue response epoch (2-5 seconds from cue-onset) for cued 10% sucrose and extinction trials (n = 9 animals, 96+ trials; two sample t-test: p = 0.909). (i) Sagittal section cartoon displaying target injection region, cell-types and fiber implant locations for mNAcSh-VTA^Pdyn^ recordings. (j) Representative coronal section showing Pdyn-SPN terminals in VTA expressing GCaMP6s underneath the fiber implant (scale bar = 150 um). (k) Mean mNAcSh-VTA^Pdyn^ reward-response for every trial of each variable sucrose condition. (l) Mean mNAcSh-VTA^Pdyn^ activity over reward response epoch (2-12 seconds from reward-onset) for each condition (n = 7 animals, 143+ trials, one-way ANOVA: 3% vs 10%, *p = 0.031). (m) Mean mNAcSh-VTA^Pdyn^ reward-response for every uncued 10% sucrose trial and every cued 10% sucrose trial. (n) Mean mNAcSh-VTA^Pdyn^ activity over reward response epoch (2-12 seconds from reward-onset) for uncued 10% sucrose, cued 10% sucrose and extinction trials (n = 7 animals, 71+ trials, one-way ANOVA: Uncued vs Ext., **p = 0.009, Cued vs Ext., ***p < 0.001). (o) Mean mNAcSh-VTA^Pdyn^ reward-response for every trial of day 7 Pavlovian extinction. (p) Mean mNAcSh-VTA^Pdyn^ activity over cue response epoch (2-5 seconds from cue-onset) for cued 10% sucrose and extinction trials (n = 7 animals, 71+ trials, two sample t-test: p = 0.387).

**Extended Data Figure 8.**
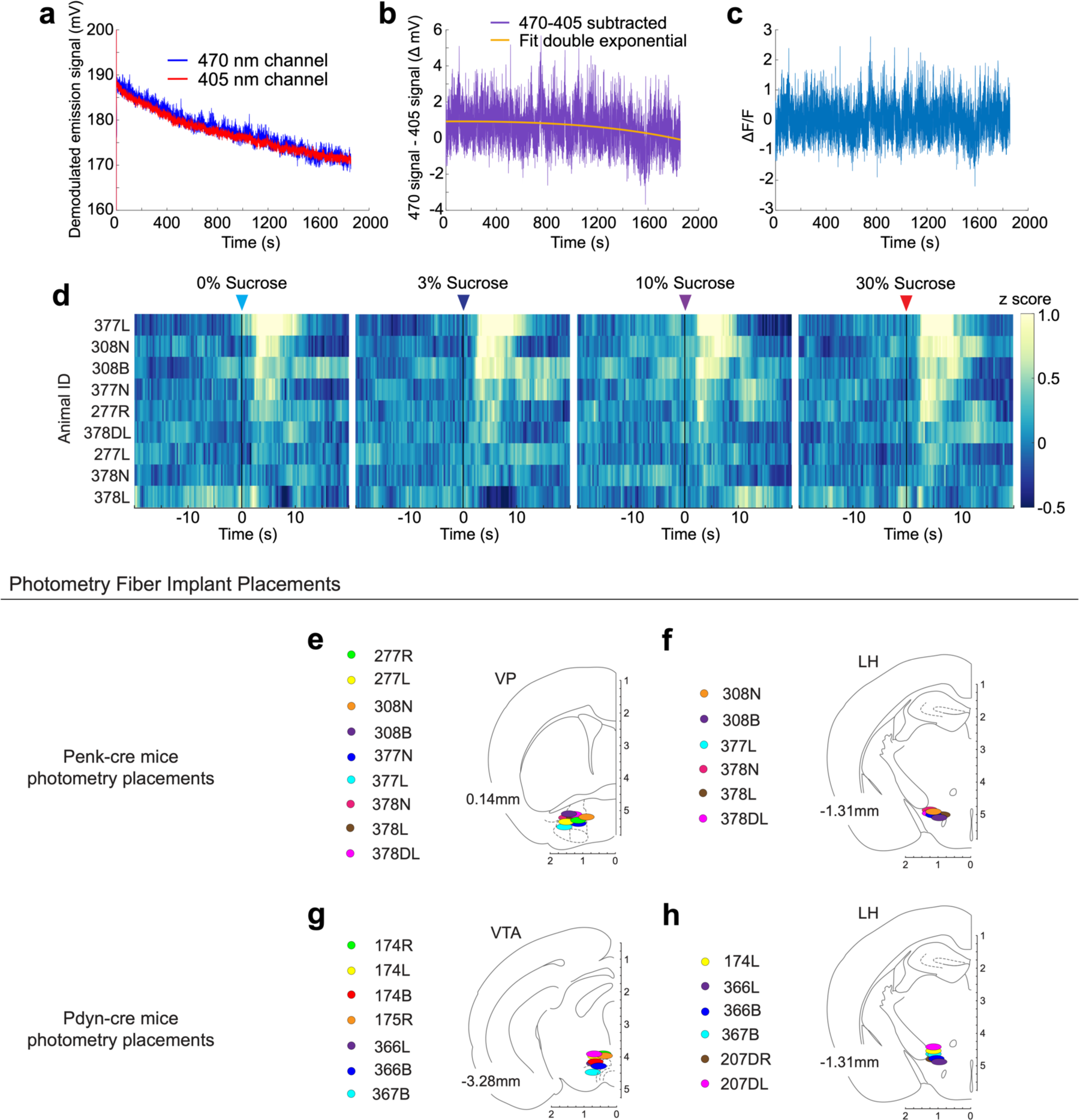
Fiber photometry signal preprocessing and implant placements. (a) Demodulated fluorescence timeseries for 405nm and 470 nm GCaMP6s excitation from same recording session. (b) Fitting a second exponential curve to the 470nm-405nm subtracted timeseries and subtracting the fitted curve to ‘detrend’ the slow drift artifact from the whole session timeseries. (c) Final normalized trace after calculation of ΔF/F using median of initial 470nm channel for division. (d) Mean mNAcSh-VP^Penk^ reward-response for 9 animals across all variable sucrose conditions. (e) Surgical placements of 400 um diameter photometry fiber implants in VP of Penk-cre animals. (f) Surgical placements of 400 um diameter photometry fiber implants in LH of Penk-cre animals. (g) Surgical placements of 400 um diameter photometry fiber implants in VTA of Pdyn-cre animals. (h) Surgical placements of 400 um diameter photometry fiber implants in LH of Pdyn-cre animals.

**Extended Data Figure 9.**
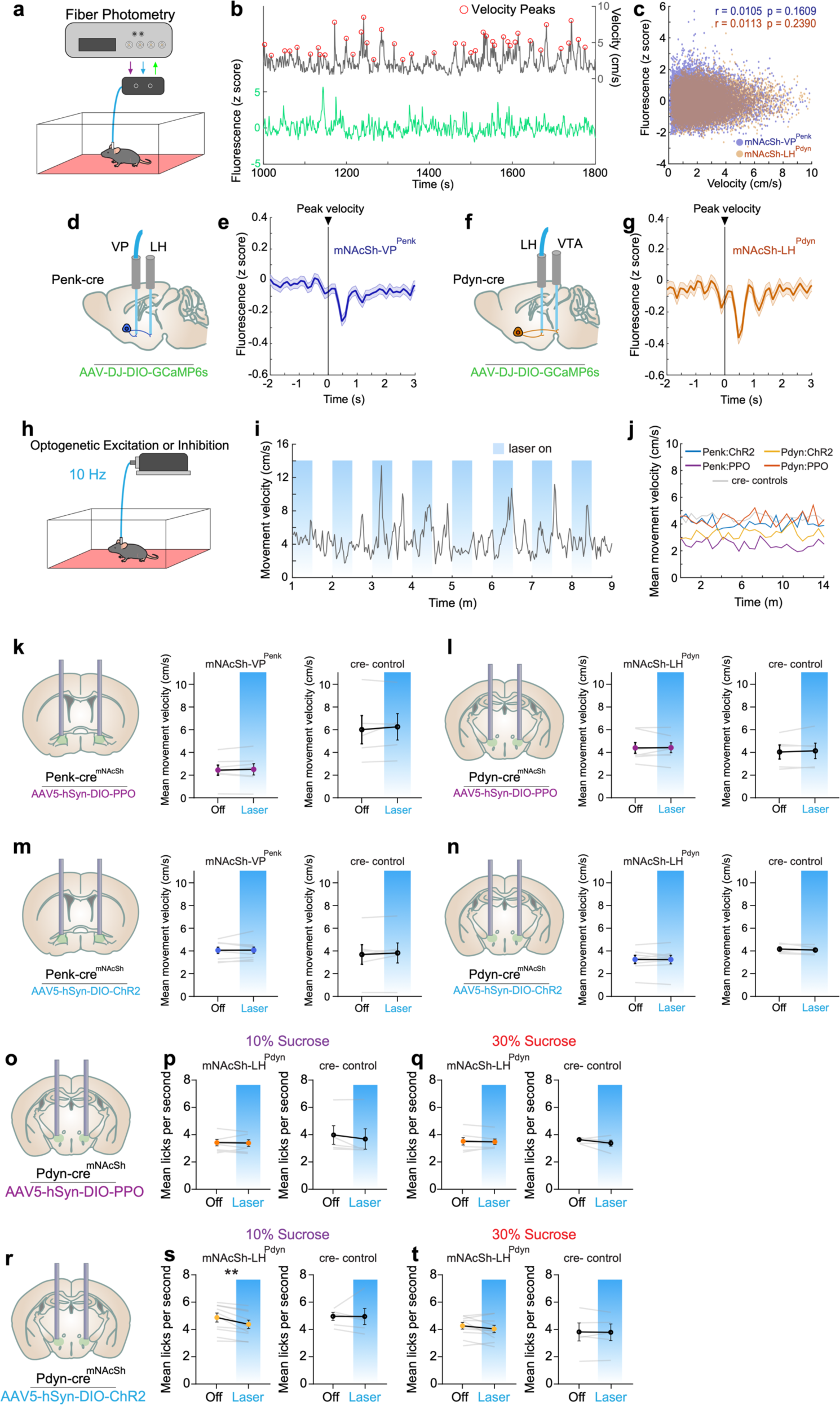
mNAcSh-VP^Penk^ and mNAcSh-LH^Pdyn^ are not recruited during locomotion and cannot disrupt locomotion. (a) Schematic of fiber photometry recording during locomotion in a behavioral arena in red light conditions. (b) Simultaneous recording of fiber photometry signal and tracking of animal movement velocity. Detection of instantaneous velocity peaks was performed post-hoc to identify movement bouts. (c) Correlation between 1 second time bins of movement velocity and 1 second time bins of GCaMP6s fluorescence (Pearson’s correlation; n = 10 Penk-cre mice, 6 Pdyn-cre mice). (d) Sagittal section cartoon displaying target injection region, cell-types and photometry recording location. (e) Mean mNAcSh-VP^Penk^ activity during every movement velocity peak. (f) Sagittal section cartoon displaying target injection region, cell-types and photometry recording location. (g) Mean mNAcSh-LH^Pdyn^ activity during every movement velocity peak. (h) Schematic of optogenetic stimulation (30 seconds on, 30 seconds off, repeating) during locomotion in a behavioral arena in red light conditions. (i) Representative timeseries of movement velocity throughout behavioral session, with alternating stimulation schedule displayed. (j) Mean movement velocity throughout duration of session for all stimulation and control conditions. (k) Within session comparison of movement velocity during mNAcSh-VP^Penk^ photo-inhibition via PPO (n = 7 treatment, 5 control animals; pairwise t-test: p = 0.497, p = 0.378). (l) Within session comparison of movement velocity during mNAcSh-LH^Pdyn^ photo-inhibition via PPO (n = 7 treatment, 5 control animals; pairwise t-test: p = 0.903, p = 0.489). (m) Within session comparison of movement velocity during mNAcSh-VP^Penk^ photo-excitation via ChR2 (n = 9 treatment, 6 control animals; pairwise t-test: p = 0.910, p = 0.440). (n) Within session comparison of movement velocity during mNAcSh-LH^Pdyn^ photo-excitation via ChR2 (n = 8 treatment, 5 control animals; pairwise t-test: p = 0.923, p = 0.480). (o) Coronal section cartoon displaying target injection region, cell-type and bilateral fiber implant location. (p) Within-session comparison of 10% sucrose consumption between trials with and without reward-paired optogenetic inhibition (n = 7 treatment, 5 control animals; pairwise t-test: p = 0.776, p = 0.187). (q) Within-session comparison of 30% sucrose consumption between trials with and without reward-paired optogenetic inhibition (n = 7 treatment, 5 control animals; pairwise t-test: p = 0.698, p = 0.220). (r) Coronal section cartoon displaying target injection region, cell-type and bilateral fiber implant location. (s) Within-session comparison of 10% sucrose consumption between trials with and without reward-paired optogenetic excitation (n = 10 treatment, 5 control animals; pairwise t-test: **p = 0.002, p = 0.973). (t) Within-session comparison of 30% sucrose consumption between trials with and without reward-paired optogenetic excitation (n = 10 treatment, 5 control animals; pairwise t-test: p = 0.174, p = 0.857).

